# Bottom-up effects of a megaherbivore alter plant growth and competition regimes, promoting vegetation heterogeneity

**DOI:** 10.1101/2025.09.13.675840

**Authors:** Hansraj Gautam, M Thanikodi, Mahesh Sankaran

**Author notes:** Corresponding author: Hansraj Gautam, Department of Biology, University of Turku, Turku 20014, Finland. Contact. Email of authors: Thanikodi M, Mahesh Sankaran.

## Abstract

Megaherbivores are known to strongly influence multiple ecological processes, but their bottom-up impacts on vegetation via nutrient redistribution remain poorly understood, particularly in mesic ecosystems. Here, we investigated how woody plant communities are influenced by large nutrient inputs in the form of dung deposited by Asia’s largest herbivore, the Asian elephant (*Elephas maximus*), in the tropical forests of southern India.
We conducted field and mesocosm experiments on woody saplings to examine three mechanisms through which dung deposition by elephants can alter plant community assembly. Specifically, we tested if elephant dung input 1) creates hotspots of plant growth, and shapes plant communities by altering 2) the negative density-dependent effects of neighborhood competition, and 3) interference competition between plant functional types differing in nutrient limitation, namely nitrogen-fixers and non-nitrogen-fixers.
Our findings show that dung deposition by elephants can generate fine-scale spatial differences in woody sapling communities by creating local growth hotspots and altering competitive interactions. We analyzed relative growth rate and final sapling size in the field experiment, and found that average-sized and large saplings receiving dung inputs were buffered against the negative density-dependent effects of neighborhood competition. In the mesocosm experiment, non-nitrogen-fixing species (which are nitrogen-limited) outcompeted nitrogen-fixers in accumulating biomass when supplied with elephant dung. These outcomes were associated with changes in their relative competitive strength which was stronger for non-nitrogen fixers under dung treatment and for nitrogen-fixers under control. Such bottom-up effects on plant growth and competition can be of substantially large magnitude, as we estimated that elephants in these forests create a total of 11000 such nutrient-rich sites /km^2^/year, with each elephant redistributing ∼130 kg Nitrogen/year through this pathway and each site receiving ∼20 g Nitrogen.

**Synthesis:** Our findings on the outcomes of sapling competition highlight the role of nutrient redistribution by megaherbivores as an underappreciated driver of species interactions that can alter plant communities at fine scales, effects that are widespread across megaherbivore habitats. Such bottom-up effects of megaherbivores, along with their top-down effects, have important conservation implications and can help in restoring species interactions and spatial heterogeneity in plant communities in defaunated habitats.

## INTRODUCTION

Large herbivores are integral to the functioning of many ecosystems as they regulate multiple ecosystem processes (Pringle et al., 2023). They have pronounced impacts on biodiversity through their effects on vegetation structure and composition (Jia et al., 2018; Van Der Plas et al., 2016), nutrient cycling (Augustine et al., 2003; Coetsee et al., 2023; Pastor et al., 2006; Villar et al., 2021) and fire regimes (Archibald & Hempson, 2016; Karp et al., 2024), in addition to downstream effects on other organisms (McCleery et al., 2024; Trepel et al., 2024). Worryingly, while ecosystems worldwide have become functionally degraded due to the body-size-downgrading of herbivore communities (Dirzo et al., 2014), our current understanding of their functional roles is strongly biased, both taxonomically and geographically (Pringle et al., 2023; Ripple et al., 2015; Sankaran & Ahrestani, 2016; Trepel et al., 2024). Such regional and taxonomic blind-spots are particularly evident for megaherbivores i.e., herbivores with >1000 kg body weight (Hyvarinen et al., 2021).

Megaherbivores are thought to have disproportionate impacts on ecosystem functioning (Owen-Smith, 1988) and are considered keystone species (Malhi et al., 2016). Yet, empirical understanding of their various ecological roles is scarce for most species (systematic review in Hyvarinen et al., 2021), which hinders a generalized understanding of the functional importance of a very large body size (Pringle et al., 2023). This limitation impedes locally informed application of such knowledge in conservation, particularly in rewilding and restoration of ecological functions lost from defaunated ecosystems (Geremia et al., 2025; Ripple et al., 2015; Svenning et al., 2016).

Among large herbivores, the disproportionate ecological impacts of a very large body size stem from its association with unique functional traits (Owen-Smith, 1988; Pringle et al., 2023). For instance, the large daily forage requirement, generalist diet and destructive foraging behaviour of megaherbivores exert strong top-down control on plants, while their wide-ranging behaviour promotes long-distance transport of nutrients and seeds (Berzaghi et al., 2018; Malhi et al., 2016; Trepel et al., 2024). Being largely immune as adults to top-down control by non-human predators, megaherbivores are instead bottom-up limited by resource availability – such bottom-up limitation amplifies their effects on vegetation (Hyvarinen et al., 2021). In recent reviews, their ecological roles have been emphasized as so uniquely important for conservation that their per-capita impact cannot be compensated for by managing densities of smaller herbivores (Hyvarinen et al., 2021; Pringle et al., 2023).

Recent meta-analyses of herbivore exclusion experiments have concluded that megafauna alter the composition of plant communities (e.g. species richness and evenness, Jia et al., 2018) and enhance spatial heterogeneity in vegetation and soil nutrients (Trepel et al., 2024). Such megafauna-induced changes in community composition and heterogeneity can arise via top-down (consumptive effects) and/or bottom-up (alteration of resource availability) regulatory pathways (Ferraro et al., 2024; Hobbs, 1996; Knapp et al., 1999; Pringle et al., 2023). Past studies on megaherbivores have predominantly focused on the top-down regulation of vegetation through consumption and trampling, with most research concentrated in African savannas (Hyvarinen et al., 2021). In contrast, their bottom-up effects are poorly understood and have been studied primarily through the lens of seed dispersal (Campos-Arceiz & Blake, 2011). A critical bottom-up impact of megaherbivores is nutrient distribution and cycling (through dung, urine and carcasses, Wolf et al., 2013), but this function remains largely understudied for megaherbivores other than hippos and savanna elephants (Hyvarinen et al., 2021). Studying the impacts of this key ecological role can shed light on the coexistence dynamics (Sitters & Olde Venterink, 2021), structure and heterogeneity of plant communities.

Megaherbivores are strong agents of nutrient cycling as they return large amounts of nutrient-rich faeces and urine (Le Roux et al., 2020; Pastor et al., 2006; Wolf et al., 2013) and facilitate long-distance transport of nutrients, like “arteries” of ecosystems (Malhi et al., 2016; Wolf et al., 2013). The patchy deposition of nutrient-rich faeces by megaherbivores creates a heterogeneous matrix of nutrient availability (Afzal & Adams, 1992; Ferraro & Lienau, 2025; Le Roux et al., 2020), which in turn, can generate fine-scale mosaics of vegetation in multiple ways. These nutrient-mediated effects could alter plant growth and survival (Tandalla et al., 2024), as well as changes in ecological interactions among plants such as competition (Van der Waal et al., 2009). For instance, herbivore dung can alter the competitive outcomes between species differing in nutrient limitation, thereby shaping community composition (Sitters & Olde Venterink, 2018, 2021). Because such responses are mediated by the quantity of nutrient inputs via dung (Sitters & Olde Venterink, 2021; Valdés-Correcher et al., 2019), megaherbivores are expected to exert disproportionate effects since they deposit large amounts of dung. However, empirical examinations of such bottom-up impacts remain scarce (Kalbitzer et al., 2019; Sitters & Olde Venterink, 2021). Therefore, our study aimed to elucidate the mechanisms through which nutrient deposition by megaherbivores influences plant communities, using elephants as a model system.

Elephants, the world’s largest herbivores, are often referred to as “ecosystem engineers” or “megagardeners” due to their large impacts on ecosystem processes (Campos-Arceiz & Blake, 2011; Poulsen et al., 2018). Existing knowledge on their ecological roles largely derives from studies on the African savanna elephant (Guldemond et al., 2017; for example, Sukumar, 2003 pp. 223-252) that occupies arid and semi-arid open habitats. In contrast, the African forest (*Loxodonta cyclotis*) and Asian elephants remain understudied, with research primarily focused on their role as seed dispersers (Campos-Arceiz & Blake, 2011; Sekar et al., 2017; but see Berzaghi et al., 2023; Blake, 2002; Omeja et al., 2014; Sivasubramaniyan & Sivaganesan, 1996). Studying these two species can elucidate the impacts of megaherbivores in wetter niches, a key gap identified in Hyvarinen et al.’s systematic review (2021).

Currently, the impacts of nutrient distribution by elephants on vegetation are poorly understood for all three species. Experimental manipulation of dung input can reveal its effects on vegetation (Kalbitzer et al., 2019; Sitters & Olde Venterink, 2021; Valdés-Correcher et al., 2019), but to our knowledge, only one such study exists. Interestingly, this study by Kalbitzer et al. (2019) did not find growth-enhancing effects of elephant dung on plants in central African rainforests. Given the lack of other field experiments, further evaluation of the supposedly large bottom-up impacts of elephants on vegetation is required.

Here, we conducted field and ex-situ mesocosm experiments manipulating elephant dung input to woody saplings in the mesic tropical forests of southern India to understand bottom-up regulation of vegetation by megaherbivores. We hypothesized that elephants create fine-scale differences in nutrient availability through dung deposition, which in turn, alters plant growth and competition regimes, with implications for local heterogeneity in vegetation (Fig 1). We tested three predictions linked to the distinct mechanistic responses by plants. First, we expected dung-deposition sites to act as growth hotspots compared to adjacent control sites (*Prediction 1*). Second, we predicted that such nutrient input via dung provides a localized buffer against negative density-dependent effects on growth (also known as the neighborhood effect or scramble competition, *Prediction 2*). We then examined how dung input affects competitive balance between plant functional types (PFT) differing in nutrient limitation (inter-PFT interference competition). To this end, we chose nitrogen-fixing (N-fixers) and non-nitrogen-fixing (non-N-fixers) woody plants; these two PFTs present an excellent contrast model as they fundamentally differ in their ability to acquire nitrogen, which often limits plant growth in many ecosystems (Vitousek & Howarth, 1991). Non-N-fixers rely solely on available forms of nitrogen in soils and thus benefit more from external nitrogen inputs, whereas N-fixers can additionally also access atmospheric di-nitrogen through symbionts (although this N-fixing ability can be expensive and is typically limited by other nutrients like Phosphorous, Tandalla et al., 2024; Varma et al., 2018). Localized deposition of nitrogen-rich dung is expected to alter the coexistence dynamics between these two plant types by alleviating nitrogen-limitation for non-N-fixers. While a previous study showed that herbivore dung alters tree-grass coexistence by shaping competition between grasses and woody N-fixers (Sitters & Olde Venterink, 2021), understanding dung-impacts on the competition between woody non-N-fixers and woody N-fixers is important in mesic ecosystems where woody non-N-fixers are more abundant. We predicted that woody non-N-fixers, being nitrogen-limited, would be able to better compete with N-fixers when supplied with nitrogen-rich dung, such that dung addition would lower the relative competitive ability of N-fixers (relative to non-N-fixers) as compared to control conditions (*Prediction 3*). Finally, to understand the overall magnitude of these bottom-up effects, we estimated the annual production of such nutrient-rich sites by elephants.

**Figure 1.**
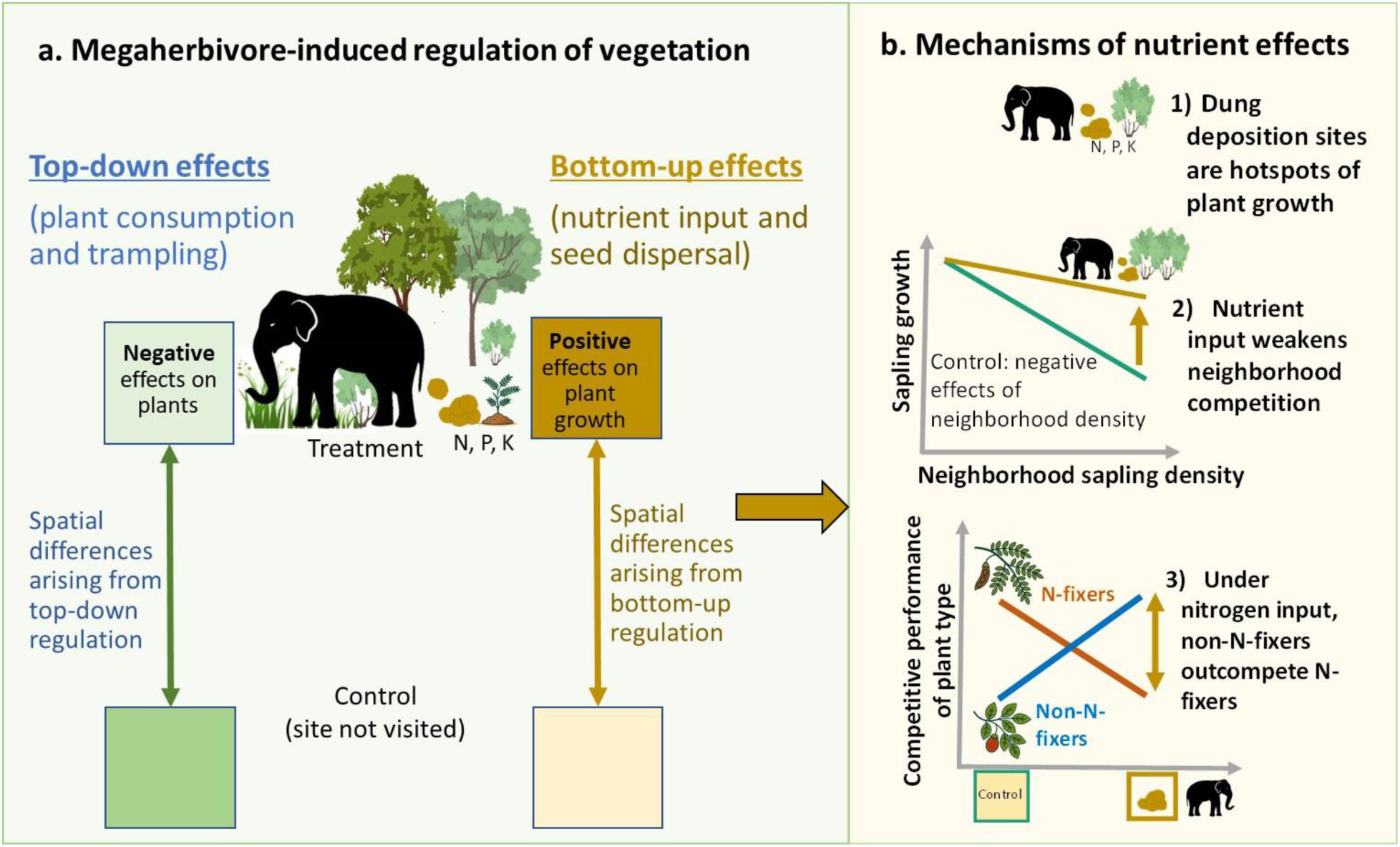
**a)** Top-down and bottom-up effects of megaherbivores as distinct regulatory pathways that can enhance local heterogeneity by promoting fine-scale spatial differences in vegetation, **b)** Hypothesized mechanisms through which nutrient input via dung deposition can create fine-scale differences between vegetation that receives such nutrient input and the adjacent control sites.

## MATERIALS AND METHODS

We conducted this study in and around the tropical forests of Nagarahole National Park (henceforth called Nagarahole). Tropical moist deciduous forests, tropical dry deciduous forests and teak plantations are the major vegetation types in the park, with mesic and hot conditions characterized by clear seasonality in rainfall and temperature (Fig. 2b). In the two decades preceding this study, mean annual rainfall ranged from ∼1000-2300 mm and ambient temperature ranged from ∼11 ℃ to ∼36 ℃ (Fig. 2a, Table S8, Supplement text B). The soil is loamy, clay-loamy and deep (Reddy et al., 2016).

**Figure 2:**
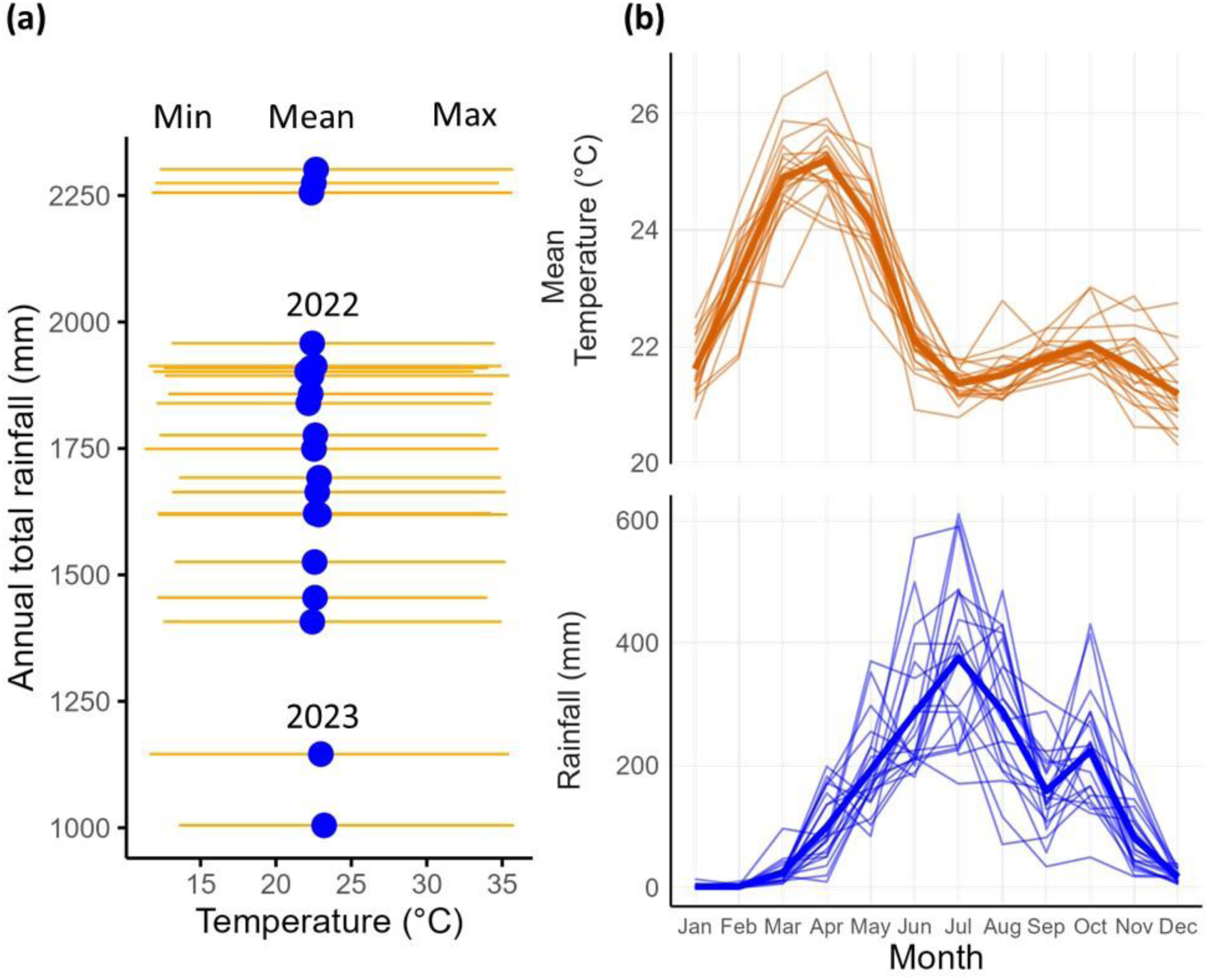
Climatic variation in Nagarahole National Park during 2012-2023 (years overlapping this study are marked). a) Inter-annual variation in total rainfall and ambient temperature (min-max range shown in orange). b) Seasonal variation in ambient temperature and total rainfall (thick lines represent long-term mean across years).

### Field experiment on woody plant growth and neighborhood competition

To examine if elephant dung deposition creates spatial differences in plant growth and neighborhood competition, we selected 19 experimental sites in September 2022 in different locations in the park, representing a range of ecological conditions spanning the three forest types. These sites were at least 2km apart from each other and were in randomly selected 2km x 2km cells, with their GPS locations pre-determined based on sampling in a previous study (*Reference removed for double-blind review*). At each location, we established two pairs of plots, 20m apart; each pair consisted of a control and a dung treatment plot (1m^2^) that were 10m apart (plot-design in Fig S1). We first sampled plots to quantify the initial state of the vegetation before applying the dung treatment. All woody individuals ≥30cm tall with diameter at breast height less than 10cm (henceforth called saplings) in plots were marked with uniquely numbered aluminum tags, and their basal diameter at 10cm aboveground measured using an INSIZE 1112-150 digital caliper (0.01mm resolution). For individuals with multiple stems, we separately measured the diameter of all stems at 10cm height, and then obtained the cumulative sum of stem diameters (henceforth called ‘basal diameter’). We also measured their height up to 1m using a scale (saplings taller than 1m were categorized into 1-2m or >2m bins).

After the initial vegetation inventory, we experimentally added dung to one plot (henceforth called treatment plot) in each pair. The treatment involved placement of an entire pile (5-10kg) of fresh dung in each plot. We collected dung piles from fresh (or overnight) defecations by adult wild elephants and transferred them to the treatment plot on the same day. We re-sampled the basal diameter and height of marked saplings after ∼6 months during April and May 2023. We also noted the status (dead/alive) of saplings; in case of saplings with multiple stems, if the main stem had died and only the smaller stem(s) survived, we classified it as ‘partially dead’. The total duration between the initial and final measurements was 162 days on average, but varied due to logistics (143-215 days). One site burned in a forest fire, leaving us with 18 sites or 36 plot-pairs.

### Ex-situ mesocosm experiment on competition between plant functional types (PFT)

To examine if dung input affected the competitive dynamics between nitrogen-fixers and non-fixers, we set up a semi-natural mesocosm experiment at a site ∼3km from the southeastern boundary of Nagarahole, in September-October 2022. The site had naturally regrowing vegetation with an overstory of deciduous trees and a dense understory. At the site, we enclosed a flat rectangular area ∼600m^2^ using a wire mesh (one-inch gap). We further covered the enclosure with a plastic net to prevent the entry of birds and mammals or falling tree branches while retaining the effects of sunlight and canopy. We selected seedlings (8-10 weeks) of native N-fixing (*Dalbergia latifolia*, *Pterocarpus marsupium*, *Butea monosperma*, *Senegalia ferruginea* (*Acacia ferruginea*)) and non-N-fixing tree species (*Terminalia tomentosa*, *Terminalia anogeissiana* (*Anogeissus latifolia*), *Tectonia grandis*, *Zizyphus oenoplia*, *Cassia fistula*, *Terminalia bellirica* and *Syzigium cumini*). These seedlings were obtained from nearby forestry nurseries (Tithimati, Beechanahalli) where seeds collected from natural forests were germinated for plantation/restoration purposes. For *Terminalia anogeissiana* and *Zizyphus oenoplia*, which are widespread in this landscape but were not available in nurseries, we collected similar sized seedlings from vegetation outside the park boundary. Together, these species represent >50% of the naturally occurring community of tree seedlings in Nagarahole forests (based on data in Gautam et al., 2019). We used a total of 132 seedlings in this experiment. Seedlings were grown in large, rectangular pots (∼18” length, 12” width, 11” depth, i.e., with surface area of 0.14 m^2^ on top) and local soil from uncultivated land adjacent to the eastern border of the park. Pebbles and plant debris in soils were first removed, after which the soil was mixed with sand in a 2:1 ratio to ensure water percolation and prevent clogging. Pots were then filled with this soil-sand mixture up to ∼90% of their heights (∼16kg), and planted with two seedlings, as described below.

We planted two seedlings (of comparable basal diameter, height) ∼10 inches apart in each pot to examine how elephant dung input influences seedling competition. We created two competition scenarios for each plant functional type (PFT): i) inter-PFT or mixture competition, i.e., competition between one N-fixer and one non-N-fixer sapling, and ii) monoculture or intraspecific competition, i.e., competition within each PFT. This resulted in two types of monocultures, one each for N-fixers and non-N-fixers. We maintained two sets of each monoculture and mixture regime pots, one being the control while the other received dung treatments, in each block (N=11 blocks of six pots containing two seedlings each). The dung treatment involved placing 500g (wet weight, ∼2/3^rd^ of a bolus) of fresh elephant dung between the two seedlings (adding whole dung pile is not possible due to limit pot size).

Since we were interested in inter-PFT competition, we ensured a balanced sample size for each PFT (N=66); however, the competition design resulted in twice the number of seedlings under monoculture than the mixture (inter-PFT) competition. Replication at the species level was limited by logistical constraints (availability of seedlings in nurseries, soil requirements for large pots, and enclosure size). At the seedling level, this experiment had a 2 (dung/control treatment) × 2 (PFT: N-fixers/non-N-fixers) × 2 (competition type: inter-PFT/ within-PFT (or intra-specific i.e., monoculture) design. All pots were watered 2-3 times a week to prevent sapling mortality, with every pot receiving similar amount of water. After the experiment ended in April 2022 (after ∼5-6 months), all saplings were harvested for dry biomass measurements. We separated shoots from roots (at the soil mark) to quantify shoot and root biomass separately, and then added them to obtain final sapling biomass. Any soil sticking to the saplings was brushed off, after which the saplings were dried in an oven at 60 °C for 60 hours and their dry weight measured in the laboratory.

### Statistical analyses of field experiment data

Our analysis of the field experiment was based on all saplings alive at the end of the experiment (N=418 saplings in 18 plot-pairs, raw mortality rates provided in Results). The treatment and control plots did not seem to differ in initial diameter of saplings (raw plot-wise means in Fig. S1). We obtained two response variables for basal diameter: relative growth rate (RGR_diameter_) and final basal diameter. RGR was calculated as [ln (final size) – ln (initial size)]/ *t*, where *t* is the duration of experiment. We analyzed RGR, which reflects biomass production per unit existing size, as it is an effective metric for quantifying growth efficiency at the seedling/sapling stage when plants are growing exponentially. As both nutrient supply and competition can modify this efficiency independent of plant size, RGR is useful in capturing performance under different ecological conditions. In contrast, final size represents the outcome of competition and nutrient input given an initial size. Therefore, these two variables inform about two complementary aspects of sapling stage, performance (RGR) and outcome (size).

We first examined if dung treatment influenced RGR_diameter_, by computing Gaussian general linear mixed effects models with maximum likelihood (*lmer* function of the R package *lmer4* Bates et al., 2015). We included treatment (dung/control), sapling density and forest type as fixed effects, while also controlling for the effect of initial basal diameter by including it as a fixed effect because RGR of smaller saplings tends to be greater. We further included the random intercepts for species and plot-pair identity to account for potential non-independence of individuals within sites and species. All continuous predictors were standardized (mean = 0, standard deviation = 1) to make their effect sizes comparable. Further, we included the three-way and two-way interactions involving dung treatment, sapling density and initial diameter, since one of our objectives was to examine if elephant dung mediates neighborhood competition which could depend on the focal sapling’s size-dependent nutrient requirements and performance. A positive effect of the interaction involving dung treatment and sapling density would suggest that dung treatment buffers saplings against negative-density dependence. We did not simplify the full three-way interaction between treatment, sapling density and size, as all these interactions with treatment were of our *apriori* interest. Upon inspection of the Gaussian GLMM (identity link), we found strong heteroscedasticity, with residual variance increasing with initial size and differing across forest type (assessed using simulated residual diagnostics, *DHARMa* package Hartig & Hartig, 2017). We therefore fitted generalized linear mixed effects models (GLMMs) with a Student t family of error distribution and a dispersion structure dependent on forest type and initial diameter, as summarized in Table 1 (using *glmmTMB* package Brooks et al., 2017). To understand the overall effect of dung treatment, we estimated the marginal means for the two treatment levels using the *emmeans* package and tested their difference using a Wald z-test on the treatment-control contrast (*emmeans* averaged over forest types). Similarly, we examined the treatment-control contrast at specified levels of density and initial size, and obtained the conditional-effects plot for the 3-way interaction using *ggpredict* function. We also present the summary table of conditional effects of the 3-way interaction for dry deciduous forest which dominated our sample. Further, since RGR takes very small values, we used predicted RGR_diameter_ to derive predicted values for two more intuitive variables, i.e., percentage change and absolute change in diameter. Percent change was calculated as 100 × (exp (*emmean* × t) −1), and absolute change as (exp (*emmean* × t) −1) × initial diameter), for t = 215 days (full duration of the experiment) (derivations in Supplement text A). These were then visualized for three discrete initial sapling sizes, i.e., average, small and large (mean initial diameter, mean − 0.5SD, mean +1SD), for treatment and density (mean and ±1SD) contrasts. For small sapling size, we chose mean − 0.5 SD because in our dataset initial diameter is right-skewed and mean – 1SD does not exist.

**Table 1.** Generalized linear mixed model specifications for different response variables. PFT: plant functional type (nitrogen-fixer/non-nitrogen-fixer).

| Response Variable | Fixed effects | Random effects | Error distribution family | Dispersion structure (in heteroscedastic models) |
| --- | --- | --- | --- | --- |
| **<u>1. Field experiment</u>** |  |  |  |  |
| **1.1. RGR<sub>diameter</sub>**<br>(mm/mm/day) | Dung × density (std) × initial diameter (std) | Species + Plot-pair | Student t | initial diameter (std) + forest type |
| **1.2. Final diameter**<br>(mm) | Dung × density (std) × initial diameter (std) | Species + Plot-pair | Student t | log (initial diameter) + forest type |
| **1.3. Final height**<br>(cm) | Dung × density (std) × ploy(initial height (std), 2) | Species + Plot-pair | Student t | initial height (std) |
| **1.4. Mortality (1/0)** | Dung | - | Binomial | - |
| **<u>2) Mesocosm experiment</u>** |  |  |  |  |
| **2.1. Final biomass**<br>(g) | Dung × PFT × Competition type + Initial size index (std) | Species + Pot ID + Block | Student t | PFT |
| **2.2. Relative Competitive Strength** (ratio based on biomass) | Dung × PFT + Initial size index ratio (std) | Species + Pot ID | Gamma (link = log) | PFT |

**Table 2:** Conditional effects from a GLMM (with Student t error distribution, and variance structure) showing final sapling basal diameter (mm) increasing with dung input, with this effect being especially evident at high density and for larger saplings. Terms of interest are marked blue and significant *P* values are marked bold.

| Conditional effects | Estimate | S.E. | CI lower | CI upper | Pr(>\|z\|) |
| --- | --- | --- | --- | --- | --- |
| **Intercept** (ref: control, DDF, mean sapling size and density) | 12.97 | 0.197 | 12.584 | 13.355 | 0.000 |
| **Treatment (Dung)** | **0.55** | 0.259 | 0.040 | 1.053 | **0.035** |
| **Sapling density** | **-0.48** | 0.188 | -0.852 | -0.116 | **0.010** |
| **Initial girth** | **13.16** | 0.287 | 12.602 | 13.725 | **0.000** |
| **Forest type – MDF** | -0.18 | 0.136 | -0.443 | 0.090 | 0.195 |
| **Forest type – Teak** | **-0.72** | 0.339 | -1.386 | -0.056 | **0.034** |
| **Treatment (Dung) × Sapling density** | **0.63** | 0.269 | 0.103 | 1.158 | **0.019** |
| **Treatment (Dung) × Initial girth** | **0.76** | 0.386 | 0.002 | 1.513 | **0.049** |
| **Sapling density × Initial girth** | -0.53 | 0.275 | -1.070 | 0.006 | 0.053 |
| **Treatment (Dung) × Sapling density × Initial girth** | 0.66 | 0.397 | -0.118 | 1.439 | 0.096 |

Similarly, we developed GLMMs for final diameter and final height, while taking into account effects of initial size by including it as a fixed effect, as specified in Table 1. In the case of sapling height, since we had measured height only up to 1m, we excluded all saplings whose initial and final height was >1m and modelled final height for this filtered subset of 308 saplings.

We analyzed sapling mortality (binary response variable) in our full dataset (N=432) using a binomial GLMM with a logit link. We also inspected net change in the count of stems (stem-count) between initial and final measurement of saplings. Net change in stem-count can result from a) natural growth trajectory where new stems sprout and grow, and b) partial mortality i.e., mortality of some of the stems. Finally, we selected the subset of saplings whose stem-count remained unchanged (N=316) and repeated the above analyses on RGR_diameter_ and final basal diameter, with models specified in Table 1, to check if the trends of growth in this subset reflected the full dataset.

### Statistical analyses of mesocosm experiment data

To examine how dung treatment affects competition between the two plant functional types (PFT), i.e., N-fixers and non-N-fixers (henceforth inter-PFT competition), we analyzed the final sapling biomass data from the mesocosm experiment in two ways. First, we tested the effect of elephant dung on final sapling biomass, using a GLMM containing the fixed effects arising from a full interaction between treatment (control/dung), PFT (N-fixer/non-N-fixer) and competition type (mixture/monoculture). To account for any variation due to initial size, initial size index (standardized, mean = 0, SD = 1) was included as a fixed effect, while species identity and block were used as random effects. The initial size index was based on an allometric equation (diameter^2^ × height). We inspected the outcomes under inter-PFT competition to infer the effect of elephant dung on the competitive ability of each PFT. We considered whether N-fixers or non-N-fixers attained greater final biomass under the dung treatment or control, when in inter-PFT competition. For this, we obtained the estimated marginal means for further comparison.

Second, we quantified the relative competitive strength (RCS) of each plant functional type (PFT), using observed sapling total biomass as a measure of performance under each competition type within each block

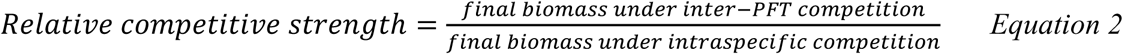

We considered a PFT to be a better competitor under inter-PFT competition if its RCS exceeded 1.0 or if it reduced the RCS of the competing PFT (Sitters & Olde Venterink, 2021). To examine this, we used a GLMM (Gamma distribution) to model RCS as a function of the fixed effects arising from a full interaction between treatment (control/dung) and PFT of the focal saplings (N-fixer/non-N-fixer). We also accounted for variation in RCS due to the fixed effects of the initial size differences between the saplings being compared (allometric size index in mixture pot/size index in monoculture pot) and the random effects of species identity. Further, we included mixture pot ID as a random effect because for the focal sapling of each PFT competing in each mixture pot (i.e. inter-PFT competition pot), two RCS values were obtained due to comparison of the focal sapling with two saplings of the same species in the corresponding monoculture pot within the block. We inferred a PFT’s competitive ability in two ways. We first tested the estimated effects of dung treatment to examine if it significantly shifted the RCS value of the focal PFT, compared to control conditions. Second, we examined the 95% confidence intervals of the estimated marginal means (using *emmeans*) of each group in the treatment × PFT interaction to test if each PFT’s RCS (95% confidence interval) was different from the null expectation of 1.0 under control and dung treatment. RCS = 1.0 implies similar performance in inter-PFT and conspecific competition. We then checked if each PFT’s “RCS>1.0 or not” status changed from control to dung treatment conditions.

### Quantifying nitrogen redistribution by elephants

To understand the broader implications of our experiments, we estimated annual rates of nitrogen redistribution per km^2^ by elephants in Nagarahole following an earlier approach to quantify carbon stocks in elephant dung (Sandhage-Hofmann et al., 2021) as follows:

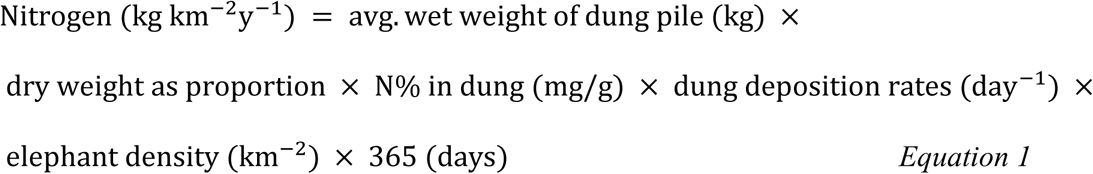

We obtained average wet weight by measuring whole fresh dung piles (N=42) in the field. To estimate dry weight, we collected ∼200-400g of sample for each dung pile, sun dried them initially, and then dried them further in the lab at 60° C for ∼60 hours. We then weighed them to obtain dry weight as a fraction of the original wet weight. Nitrogen content in elephant dung was quantified for three seasons using LECO CN analyzer (LECO Corporation, St. Joseph, MI) in a parallel study on elephant diet (*Reference removed for double-blind review*), to which we added data from another season (Table S7). Elephant density in Nagarahole and dung deposition rates of Asian elephants were taken from previous studies (Table S7).

## RESULTS

### 1. Field experiment

#### 1a) Elephant dung increases sapling growth and alters neighborhood competition

Spatial differences in sapling growth in response to dung deposition were clearly evident, supporting *Prediction 1*. The raw percentage increase in the basal diameter of saplings under dung treatment was 16% (n=194 saplings, SD=33.7), while it was only 9% in the control plots 10m away (n=224, SD=25.5; plot-wise means in Fig. S1). Accordingly, the GLMM of relative growth rate (RGR) showed that dung addition enhanced RGR_diameter_ by over 70% relative to control saplings (comparison of estimated marginal means, *P*=0.040, Fig. 3a). Density had a negative effect on the RGR_diameter_ of larger saplings under control conditions (*P* = 0.033) and while dung treatment appeared to reduce this negative effect of density on large saplings, this 3-way interaction effect was statistically not significant (*P* = 0.12; Table S1). We obtained the estimated marginal means at different sapling densities for three discrete initial sizes and found a size-dependent buffering against the negative effects of high density, consistent with *Prediction 2*. Dung treatment provided a significant advantage at average and high sapling densities for average-sized (mean initial diameter) and large (mean + 1SD) saplings, whereas this advantage was not seen at lower densities and for small saplings (Fig. 3b and 3c). Fig 3c shows these effects in terms of percentage and absolute growth in diameter derived from predicted RGR_diameter_ for discrete values of sapling density and initial size.

**Figure 3:**
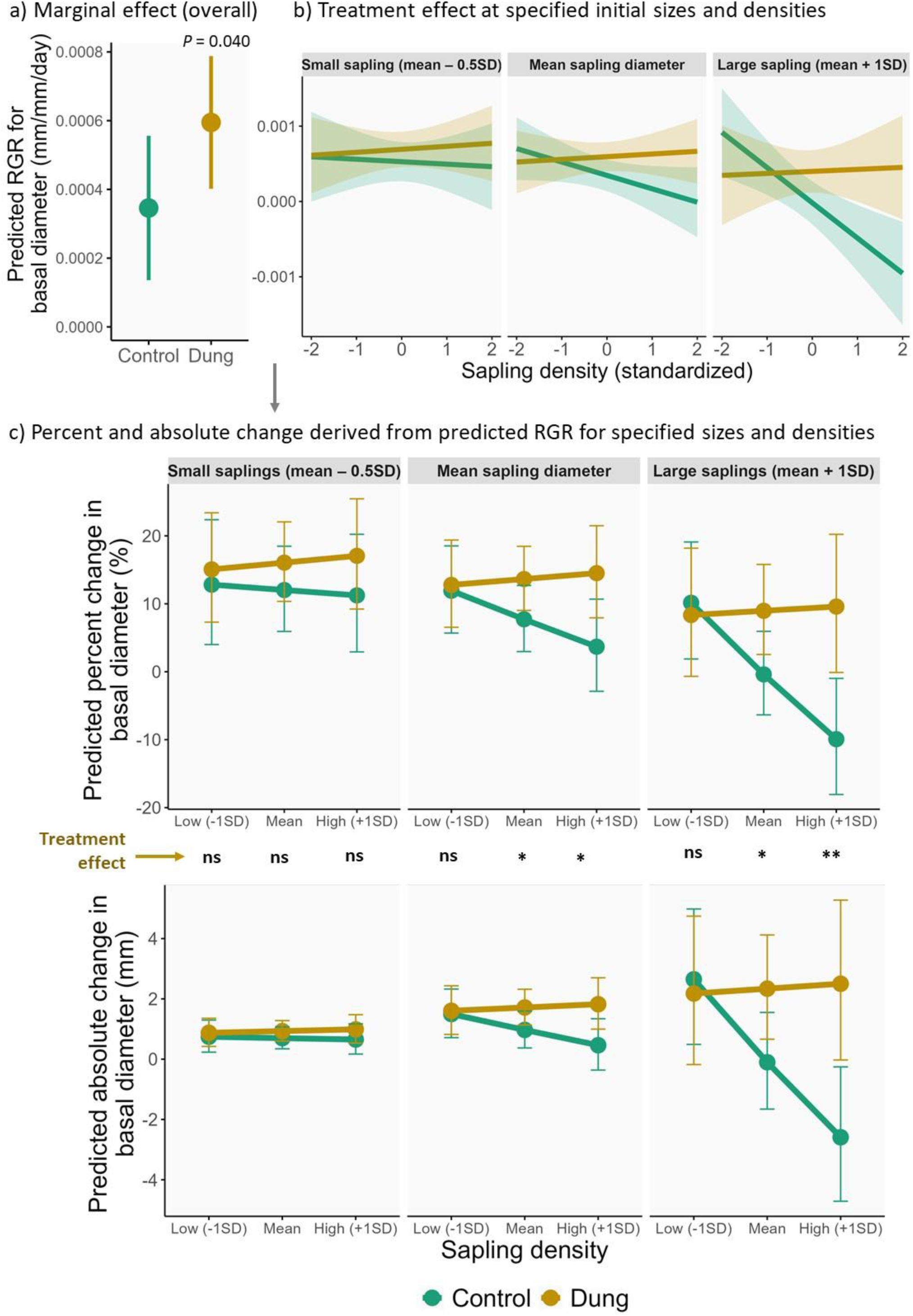
Dung treatment buffers larger saplings against the negative effects of sapling density on growth. a) Overall effect of dung treatment on RGR_diameter_ (based on estimated marginal means or *emmeans* across densities, sizes and forest types), b) Predicted RGR_diameter_ based on *emmeans* at specified densities for three discrete initial sizes (obtained using *emmeans*, raw data plotted in Fig S2), and b) The same effects expressed as percent change and absolute change in basal diameter, derived from the *emmeans* of RGR_diameter_ at specified densities and initial sizes (see Supplement text A). Error bars represent 95% confidence intervals. The discrete value of initial diameter for small sapling is defined as mean – 0.5 SD instead of mean – 1SD which is non-existent in our dataset due to its right skew. Significance codes correspond to tests of treatment-control contrasts in the predicted RGR at specified density/size (based on *emmeans*): ns: p>0.05; *: p <0.05; **: p <0.01; ***: p<0.001.

The positive effects of dung treatment were statistically more evident in the GLMMs of final sapling size (Table 1). In addition to the overall positive marginal effect of dung treatment (comparison of *emmeans*: P=0.035, Fig 4a), this effect was greater for large saplings which were also buffered against the negative effects of high sapling density under dung treatment, consistent with *Prediction 2* (Fig 4b, Table 1). In contrast, the GLMM of final height revealed that the positive marginal effect of dung treatment was statistically not significant (based on *emmeans*, *P*=0.063, Fig 4c). Saplings increased their height under crowding, but this was not influenced by dung treatment (Table S2).

**Figure 4.**
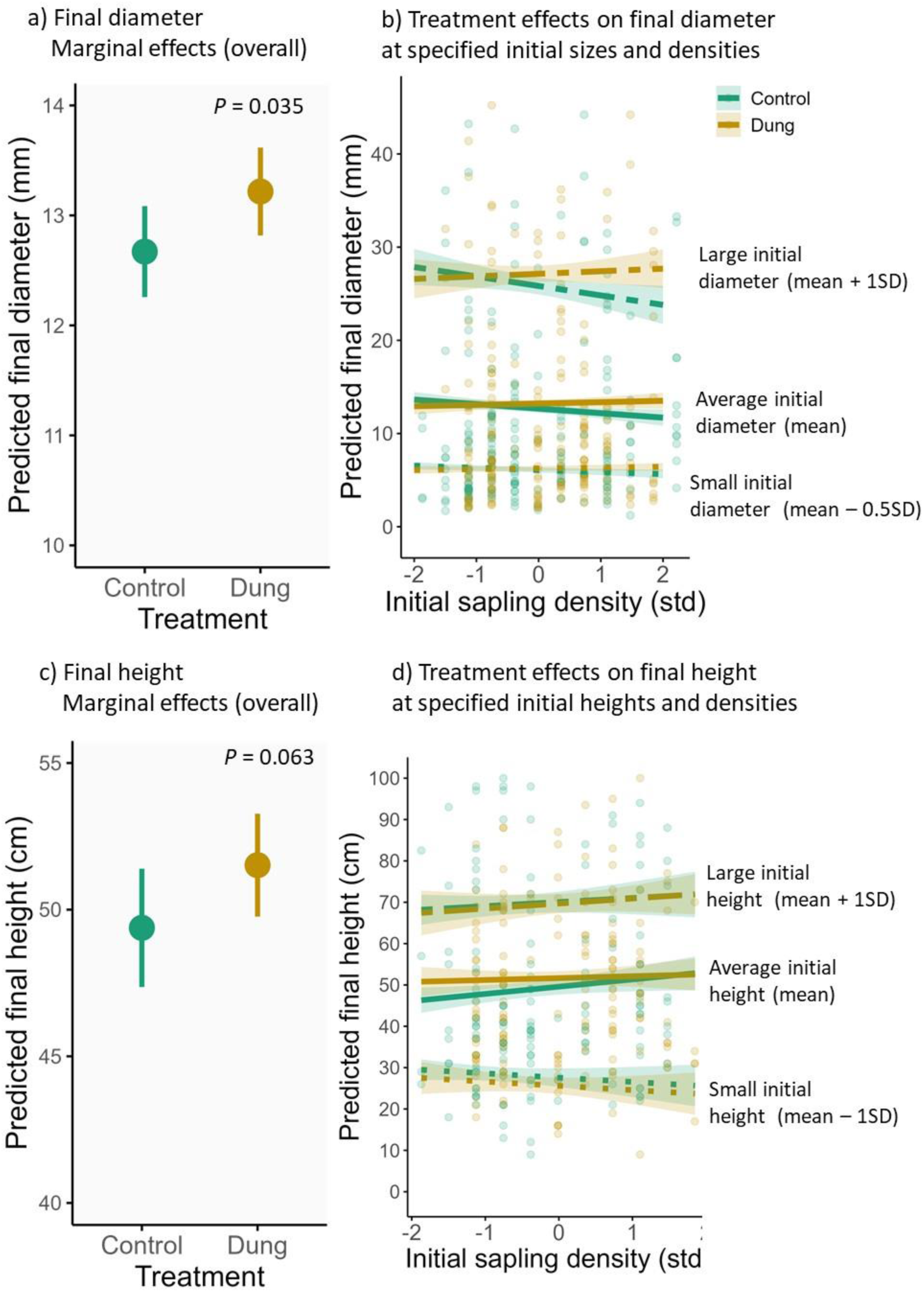
**a & c.** Estimated marginal means of final sapling diameter and final height for saplings under dung treatment and control (across the distribution of sizes, densities and forest types). **b & d.** Estimated marginal means at specified densities for discrete initial sizes, under dung treatment and control. Predicted values are shown for three discrete initial sizes. In panel b, small sapling diameter is specified as mean – 0.5 SD initial diameter instead of mean – 1SD which is non-existent in our dataset due to its right skew.

#### 1b) Sapling mortality and change in stem-count

Mortality during the ∼6-month long field experiment was low in the full dataset, with mortality rate being 3.96% under the control (8/202) and 2.61% under the dung treatment (6/230), although a binomial GLMM revealed that these differences were not significant (*P*=0.43). We also inspected the net change in stem-count for the filtered dataset containing only the saplings alive at the end of the experiment (N=418). We found that a net increase in stem-count was more common in saplings receiving the dung treatment (13.4%) than control saplings (8.8%), whereas a net loss of stem-count was marginally more common in control saplings (14.4% vs 12.1% for dung treatment, Figure S4, Supplement text A).

We selected the 76% saplings whose stem-count remained the same between the initial and final measurement (N=316), and repeated our analyses of the effects of dung treatment on RGR_diameter_ and final basal diameter for this subset, with models specified in Table 1. In this smaller dataset, while the directions of effects were broadly consistent with the findings in the full dataset, many effects were statistically not significant (Supplement Table S3 and Table S4). Nevertheless, for both RGR_diameter_ and final diameter, we found clear statistical support for dung input providing a buffer against the negative effects of density for average-sized saplings (Fig. S3).

### 2. Ex-situ mesocosm experiment

#### Non-N-fixers outcompete N-fixers under dung input

Both analyses of final sapling biomass and relative competitive strength (RCS) suggested that elephant dung affected competition between plant functional types (inter-PFT competition), supporting *Prediction 3*. When in inter-PFT competition, there was no difference in the biomass of N-fixers and non-N-fixers under control conditions (*P* = 0.178 for *emmeans* contrast, Figure 5a, Table S6b). In comparison, dung addition resulted in non-nitrogen fixers accumulating over twice the biomass as nitrogen-fixers (*emmeans* contrast: *P*=0.003).

**Figure 5:**
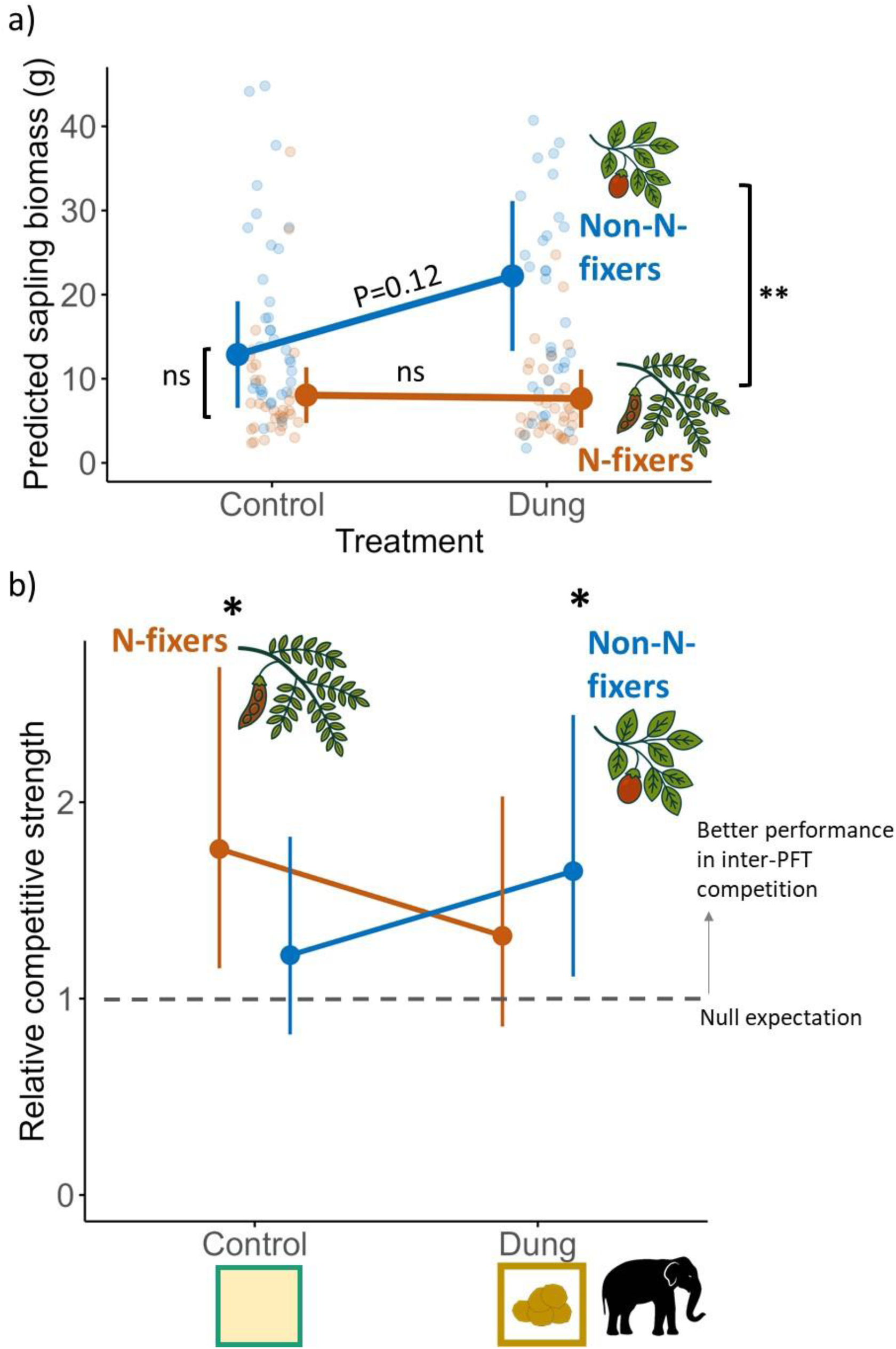
Outcomes of competition between the plant functional types N-fixers and non-N-fixers in the mesocosm experiment. **a)** Predicted total sapling biomass under competition between plant functional types (based on *ggpredict*): when supplied with dung, non-N-fixers accumulate greater biomass than N-fixers under mixture competition. **b)** Predicted relative competitive strength (RCS, based on *emmeans*): RCS is significantly >1.0 for N-fixers under control and for non-N-fixers under dung treatment (raw data plotted in Fig. S4). RCS = biomass under inter-PFT competition / intraspecific competition. Error bars represent 95% confidence intervals around estimated means.

GLMM of relative competitive strength (RCS) revealed that the dung treatment substantially increased the RCS of non-N-fixers (i.e., improved performance under inter-PFT vis-à-vis intraspecific competition) relative to control conditions, after accounting for initial size and random effects (Figure 5b, Table 3b). Dung treatment appeared to reduce the RCS of N-fixers, but not significantly so. Next, we inspected the 95% confidence intervals to test for differences in the competitor status of the two PFTs with respect to the null expectation of 1.0. In the absence of dung input, N-fixers performed better under inter-PFT competition than under intra-specific competition (mean RCS= 1.762, *P*=0.009, null RCS=1.0) (Figure 5b, Table 3a). In contrast, this better performance of N-fixers under inter-PFT competition did not hold when dung was added (RCS did not differ from 1.0). Contrastingly, the non-N-fixers showed better performance under inter-PFT competition under the dung treatment (mean RCS =1.649, *P*=0.013), which was not the case under control conditions.

**Table 3:**
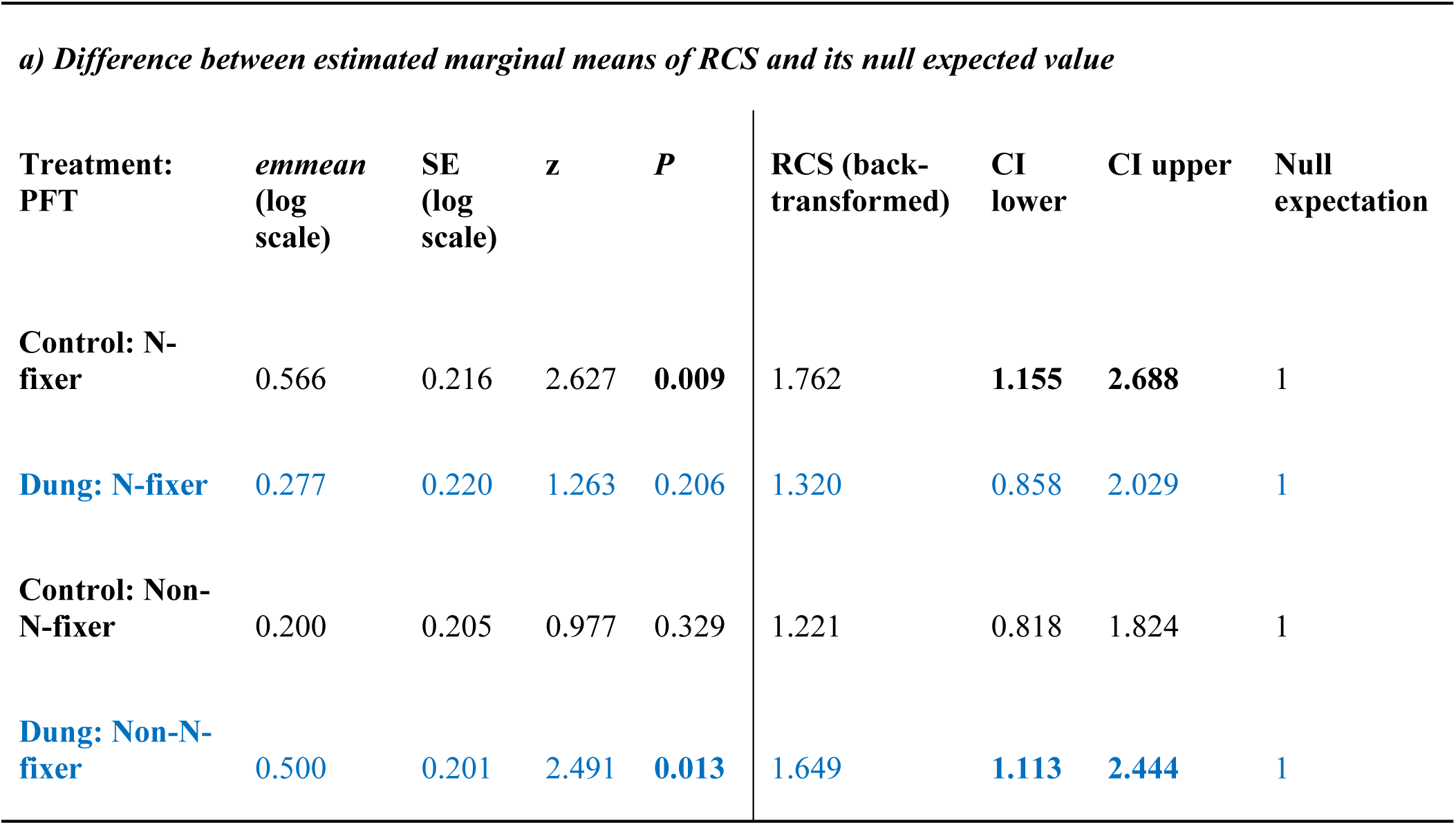
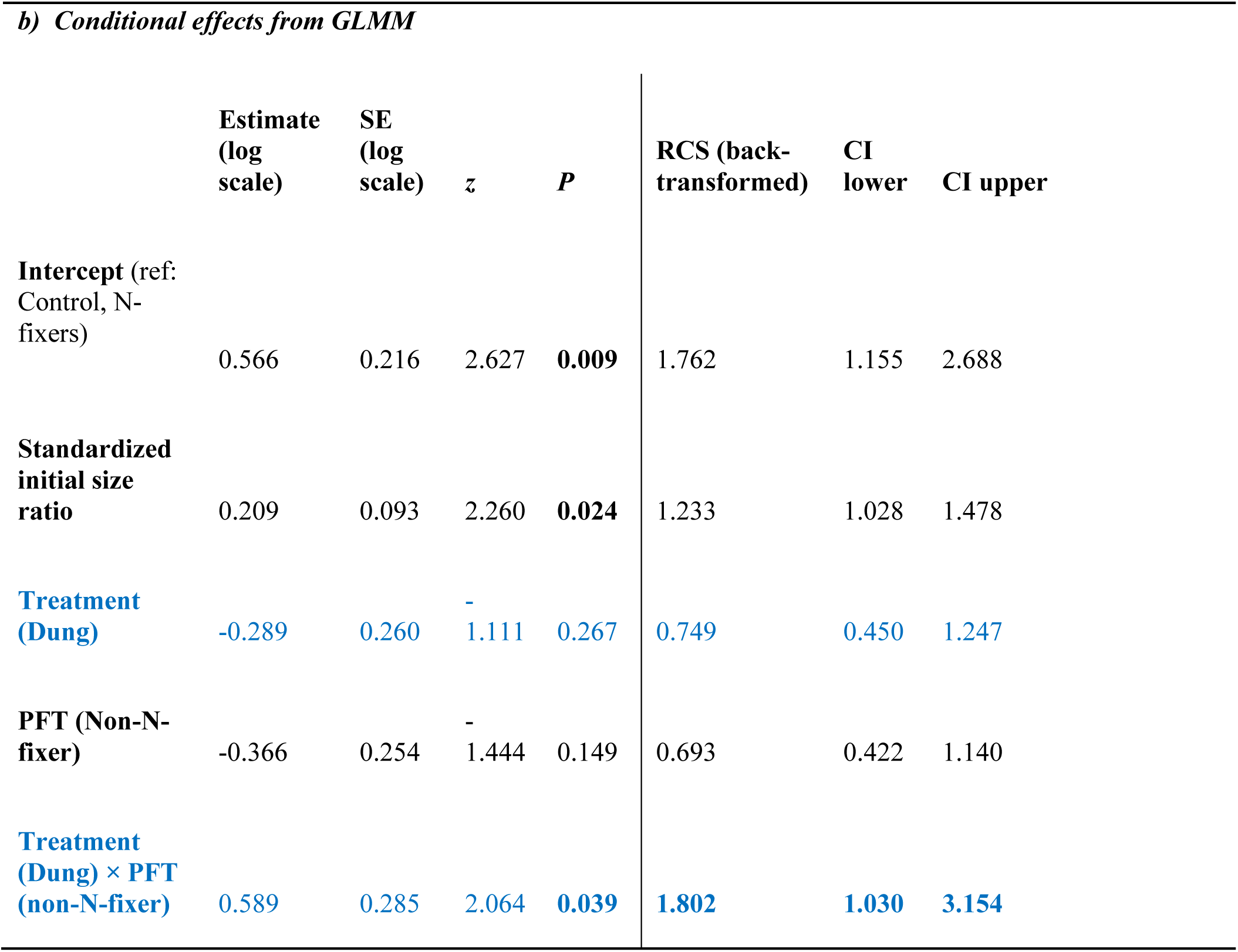
Results from the GLMM of relative competitive strength (RCS). **a)** Estimated mean RCS: N-fixers were more competitive (RCS>1.0) under while non-N-fixers were more competitive under dung treatment, **b)** Conditional effects form GLMM: non-N-fixers increased their RCS under dung treatment relative to control (rows of interest are marked in blue). Rows with P<0.05 are marked in bold.

| Treatment:<br>PFT | *emmean*<br>(log<br>scale) | SE<br>(log<br>scale) | *z* | *P* | RCS (back-<br>transformed) | CI<br>lower | CI upper | Null<br>expectation |
| --- | --- | --- | --- | --- | --- | --- | --- | --- |
| Control: N-fixer | 0.566 | 0.216 | 2.627 | **0.009** | 1.762 | **1.155** | **2.688** | 1 |
| Dung: N-fixer | 0.277 | 0.220 | 1.263 | 0.206 | 1.320 | 0.858 | 2.029 | 1 |
| Control: Non-N-fixer | 0.200 | 0.205 | 0.977 | 0.329 | 1.221 | 0.818 | 1.824 | 1 |
| Dung: Non-N-fixer | 0.500 | 0.201 | 2.491 | **0.013** | 1.649 | **1.113** | **2.444** | 1 |

*b) Conditional effects from GLMM*
|  | Estimate<br>(log<br>scale) | SE<br>(log<br>scale) | *z* | *P* | RCS (back-<br>transformed) | CI<br>lower | CI upper |
| --- | --- | --- | --- | --- | --- | --- | --- |
| **Intercept** (ref:<br>Control, N-fixers) | 0.566 | 0.216 | 2.627 | **0.009** | 1.762 | 1.155 | 2.688 |
| **Standardized**<br>**initial size**<br>**ratio** | 0.209 | 0.093 | 2.260 | **0.024** | 1.233 | 1.028 | 1.478 |
| **Treatment**<br>**(Dung)** | -0.289 | 0.260 | -<br>1.111 | 0.267 | 0.749 | 0.450 | 1.247 |
| **PFT (Non-N-**<br>**fixer)** | -0.366 | 0.254 | -<br>1.444 | 0.149 | 0.693 | 0.422 | 1.140 |
| **Treatment**<br>**(Dung) × PFT**<br>**(non-N-fixer)** | 0.589 | 0.285 | 2.064 | **0.039** | **1.802** | **1.030** | **3.154** |

### 3. The overall magnitude of nitrogen redistribution by elephants

We estimated that elephants annually produce 11038 such nutrient-rich patches/ km^2^ on average in these tropical forests, amounting to an annual cycling of 218.88 kg N/km^2^ through this pathway alone (Table S6). On a per capita basis, each adult elephant redistributes ∼130.29 kg N/year throughout its home range by creating 6570 dung sites/year (Table S6). Elephant dung piles are fertility hotspots as their nitrogen content (annual average = 0.971%) is substantially higher than the upper soil layers in this landscape’s deciduous forests (0.14 to 0.31%; Mani et al., 2018, 0.40% in Swamy and Proctor, 1994; 0.26% in Ramachandra et al., 2012).

## DISCUSSION

Due to their exceptional body size and wide-ranging behaviours, megaherbivores are expected to have a profound role in ecosystem functioning through both top-down and bottom-up impacts. Our experiments offer important insights into their bottom-up effects via large-scale redistribution of nutrients and its associated impacts on woody plant communities in mesic ecosystems. We found that elephant dung deposition sites 1) act as hotspots of plant growth (Fig 3a, 4a), 2) weaken the negative effects of high sapling density on average-sized and large saplings (Fig 3c, 4b), and 3) shift the competitive balance between plant functional types differing in nutrient limitation, by favoring non-N-fixers over N-fixers (Fig 5).

Importantly, 4) the magnitude of such effects is substantial; we estimate that elephants create >11000 nutrient-rich sites annually in these mesic tropical forests (Table S7). These findings highlight the role of megaherbivores as modifiers of species interactions key to plant community assembly, and as promoters of fine-scale vegetation heterogeneity across their vast home ranges.

### Elephant dung deposition creates hotspots of plant growth

We found that the large inputs of nitrogen (∼19.83g N, Table S7) make elephant dung deposition sites hotspots for plant growth. Elephant dung strongly enhanced growth in the diameter of woody saplings compared to adjacent control plots. However, this growth-enhancing effect was not statistically significant for plant height (*P*=0.063), possibly because height responses may take longer to manifest than diameter (eg. Noyer et al., 2019; Seymour et al., 2022), potentially reflecting preferential allocation to bark thickening before height as a strategy to reduce mortality (eg. from fire Lawes et al., 2011). Such potentially delayed effects on height could explain why the only previous experiment of this nature could not detect growth-enhancing effects of elephant dung in central Africa’s rainforests (Kalbitzer et al., 2019). Kalbitzer et al. (2019) analyzed height, and not diameter, and did not find growth-enhancing effects of elephant dung one year after experimental dung input. However, shade-tolerant saplings in their study enhanced leaf production, suggesting that the allocation of this productivity to increases in shoot length may be delayed but potentially detectable with longer monitoring. Our inspection of stem-count data suggests that the observed increase in diameter partly ensued from a net increase in stem-count in saplings receiving elephant dung input (Fig S3). Even for saplings whose stem-count remained unchanged, basal diameter appears to have grown under elephant dung treatment, although such effects were not statistically significant in this down-sized dataset. Furthermore, sapling survival appears to be enhanced by dung treatment (2.61% mortality vs. 3.96% mortality in control saplings) in our dataset, although this difference was not statistically significant, whereas Kalbitzer et al. (2019) reported significantly lower mortality under dung treatment.

### Elephants modify ecological interactions key to plant community assembly

We found that nutrient inputs via dung can locally alter plant competition in two ways. First, elephant dung can buffer saplings against neighborhood competition or negative density-dependence. This was especially evident in the case of average-sized and large saplings whose growth under control conditions declined sharply with increasing sapling density (Fig. 3-4). In contrast, dung addition was associated with a near consistent ecological release from neighborhood competition for all sapling sizes (see flattish slopes in Fig 3, Fig 4b). Since larger saplings have greater nutrient requirements, their dung-mediated release from neighborhood competition suggests that nutrient inputs from dung can be substantial. In contrast, dung input did not influence the effects of density on sapling height; increased vertical growth was only observed in response to increased sapling density, which is a well-known response to crowding, possibly mediated by light competition (Gruntman et al., 2017).

Second, our mesocosm experiment revealed that dung can modify the outcome of interference competition between plant functional types differing in nutrient limitation. Non-N-fixers accumulated more biomass and outcompeted N-fixers when amended with dung but not otherwise. That elephant dung favored non-N-fixers was also clearly evident in the analyses of relative competitive strength (RCS). The otherwise better performance of N-fixers in control conditions (RCS>1.0) did not hold in the presence of dung. Dung input implied net gains for non-N-fixers in two ways: by enhancing the competitive status of non-N-fixers (RCS > 1.0) and by lowering it for N-fixers which had high RCS in control conditions. Herbivore dung has been previously shown to alter the competitive balance between grasses and woody N-fixers in African savannas (Sitters & Olde Venterink, 2021). Taken together, these findings suggest that such bottom-up effects of herbivores can be critical in regulating the abundance of different plant types, including woody N-fixers, non-fixers and grasses.

Dung-mediated differences in local competitive outcomes can determine which saplings succeed in transitioning to the recruitment stage, thereby influencing community composition. Our findings indicate that dung deposition can locally accelerate competitive exclusion along at least two axes: 1) size and 2) nutrient limitation among plant functional types (PFTs). First, although larger saplings were negatively affected by neighborhood density under control conditions, they were buffered against neighborhood competition in the presence of dung (Fig. 3, Fig. 4). This similarity in RGR between large and small saplings, especially at high densities (Fig. S2), suggests that herbivores can reinforce or even amplify existing size differences through their dung inputs (Fig. 4). Such dung-mediated ecological release from competition can reinforce the beginners’ advantage for larger saplings (for example, taller understory plants have an advantage in light competition), thus potentially excluding smaller saplings. Second, along the nutrient-limitation axis, herbivore dung addition is likely to favor nutrient-limited plant species. In our mesocosm experiment, nitrogen-rich dung shifted the competitive balance between non-N-fixers and N-fixers (Fig 4), suggesting that dung deposition could bring about fine-scale spatial differences in the relative abundances of these two plant types by favoring non-N-fixers. It must be noted that the observed effects on inter-PFT competition in our mesocosm experiment may be an underestimate as we used only <10% (500g) of the amount of an average elephant defecation (6.88 kg) in our treatments. Importantly, such effects on the coexistence among plant species may be widespread along other axes of nutrient limitation, for example, fast- vs- slow-growing and shade-tolerant vs. light-demanding species. Localized competitive exclusion along these axes could thus change plant communities in multiple ways. Such dung-mediated gains by large, nutrient-limited plants at the expense of smaller, non-nutrient-limited saplings can potentially contribute to the decline in tree community richness often reported in herbivore presence (Jia et al., 2018).

### Conservation implications

Our study has implications for understanding how defaunation may impact ecosystem functioning through effects on species interactions shaping plant communities. Poulsen et al. (2018) speculated that the loss of nutrient cycling associated with megaherbivores would result in the loss of light-demanding species as they are more nutrient-limited. While we did not focus on the light-demanding/shade-tolerant distinction, elephant dung indeed favored nutrient-limited plant types in our experiment (non-N-fixers) and thus elephant presence can substantially shift the composition of plant communities, possibly even at the landscape scale through cumulative generation of numerous temporary growth hotspots over multiple years. Nevertheless, we note the contradictory earlier findings of Kalbitzer et al. (2019) where elephant dung had small positive effects on shade-tolerant species that are less nutrient limited, although these results were based only on leaf production. In addition to the effects at the landscape level, we propose that enhanced fine-scale heterogeneity in community composition is another key consequence of such nutrient redistribution by megaherbivores.

The large-scale annual production of nutrient hotspots by elephants (totally >11000/km^2^/year in Nagarahole) likely amplifies vegetation heterogeneity through fine-scale differences in growth and competition, which is in line with earlier findings that herbivore dung deposition increases soil heterogeneity (Afzal & Adams, 1992; Ferraro & Lienau, 2025; Hobbs, 1996). Importantly, the functional importance of megaherbivores is underscored by the fact that the amount of dung input is a key modulator of vegetation responses (Valdés-Correcher et al., 2019), and this input is exceptionally large in the case of megaherbivores. It is likely that such oversized nutrient-mediated impacts could also have downstream effects on other organisms. Thus, protecting megaherbivore populations is critical to conserving such important functional roles.

### Future directions

Our experiments open several possibilities to further understand bottom-up effects of large herbivores. As these dung sites are temporary nutrient hotspots, they would only induce short-term spatial differences in plant growth. To what extent these pulses of altered growth trajectories shape the recruitment fate to mature stages can only be understood through longer-term studies, or models that consider such pulsed nutrient effects on plant growth and competition. Second, we encourage investigations on competition between plant functional types along other axes of differentiation, including fast- vs slow-growth and shade-tolerant vs light-demanding species. Third, stoichiometric ratios of different growth-limiting elements in dung can be important for plant growth and interactions. Megaherbivores have high phosphorous requirements and thus their faeces tend to be P-depleted i.e., with high N:P ratios (Le Roux et al., 2020) that could favour non-N-fixers over N-fixers (Sitters & Olde Venterink, 2021).

In conclusion, our study highlights large-scale nutrient redistribution by megaherbivores as an underappreciated driver of plant growth and species interactions that generate fine-scale vegetation heterogeneity. Such functional roles of megafauna remain understudied in the mesic tropics (Hyvarinen et al., 2021), particularly in South and Southeast Asia (Sankaran & Ahrestani, 2016). These regions have faced extensive defaunation and deforestation in both modern and prehistoric times (Svenning et al., 2016). Localized restoration efforts in such ecosystems could benefit from a mechanistic understanding of herbivore impacts (see Geremia et al., 2025), including the relative impacts of top-down and bottom-up regulation. Our findings suggest that large herbivore rewilding could benefit conservation actions by restoring species interactions and fine-scale vegetation heterogeneity, although such ecological roles should be seen more as an outcome of well-preserved ecosystems. We call for further investigations to advance a more comprehensive understanding of herbivore functions in these understudied ecosystems.

## ACKNOWLEDGEMENTS

We are grateful to the Principal Chief Conservator of Forests, Karnataka Forest Department, the office of the Conservator of Forests and Director, Rajiv Gandhi (Nagarahole) National Park, Hunsur, for field research permits and other forest officials for facilitating our fieldwork. We are also grateful to the Manager, Jungle Lodges and Resorts, Kabini, for providing us the space to conduct the ex-situ experiment, and to the forestry nurseries in Tithimati and Beechanahalli for providing us seedlings. We especially thank Prof. T.N.C. Vidya and the Kabini Elephant Project for logistical support during fieldwork as well as for the collaboration under which the fieldwork was carried out. We are thankful to our field assistants Krishna, Pramod and Shankar for assisting with fieldwork and ensuring our safety. We thank Abhirami Ravichandran for help during nutrient analyses of fecal samples. We are thankful to the two reviewers for the very helpful critical feedback on the first draft and to Mayank Kohli for feedback on a pre-submission draft. HG analyzed and wrote this paper while working at the University of Turku.

## AUTHOR CONTRIBUTIONS

HG: funding acquisition, conceptualization and research design, fieldwork and laboratory analyses, formal statistical analyses, writing the first draft, review and editing; MT: fieldwork; MS: research design, review and editing. All authors read and approved the manuscript for submission.

## Data Availability Statement

All data files and scripts used to prepare plots and analyses tables are attached as supplementary files. The data will be uploaded on a publicly accessible repository platform (Zenodo) after acceptance.

## SUPPLEMENTARY INFORMATION

**Figure S1:**
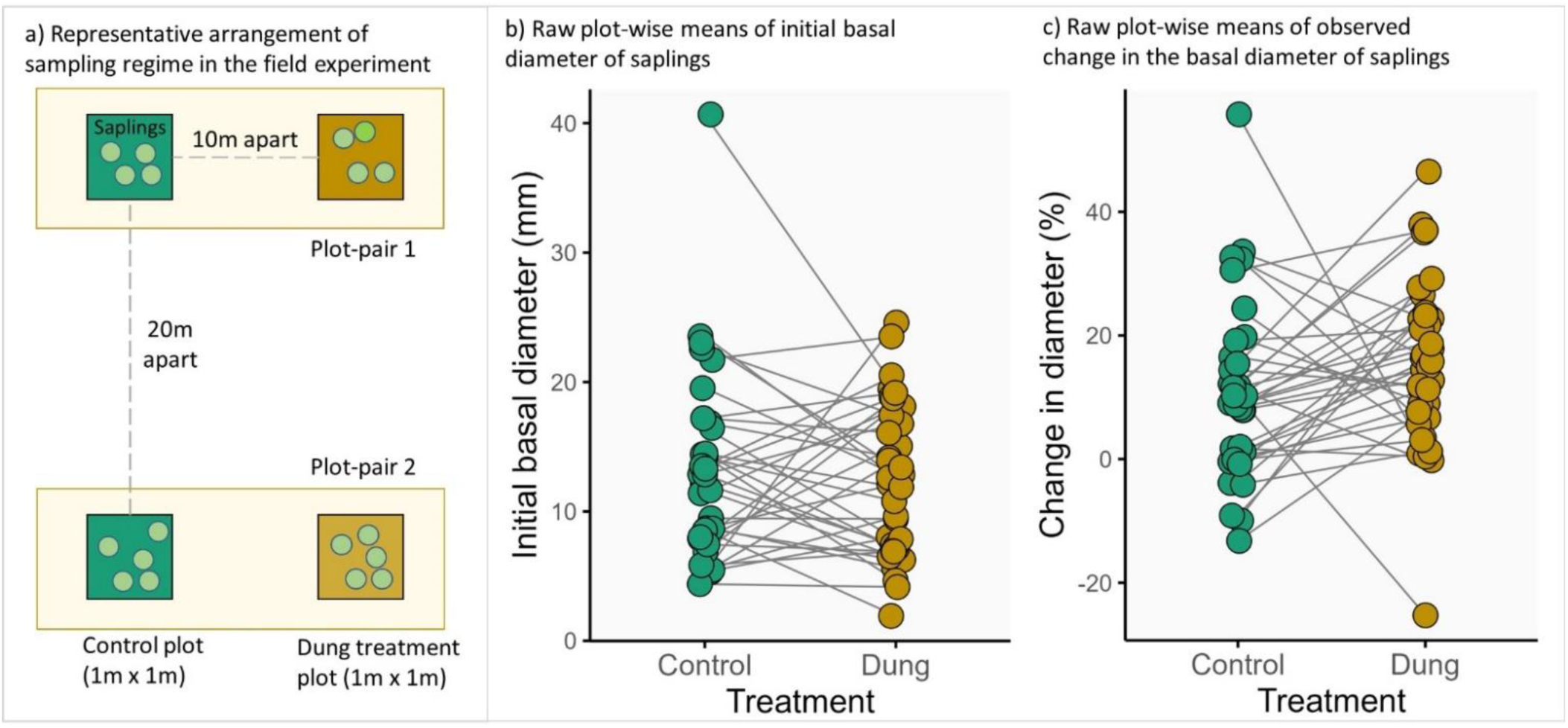
a) Representative arrangement of the sampling regime in the field experiment on sapling response to elephant dung treatment. b) Raw plot-wise means of initial basal diameter of saplings in the plot pairs (averaged across all individuals within each plot), c) Raw plot-wise means of observed percentage change in the basal diameter of saplings in 36 plot-pairs during the course of the field experiment (i.e., change between initial and final measurement). An increased growth in response to dung treatment is evident in the majority of plot pairs, relative to control plots 10m away.

**Figure S2.**
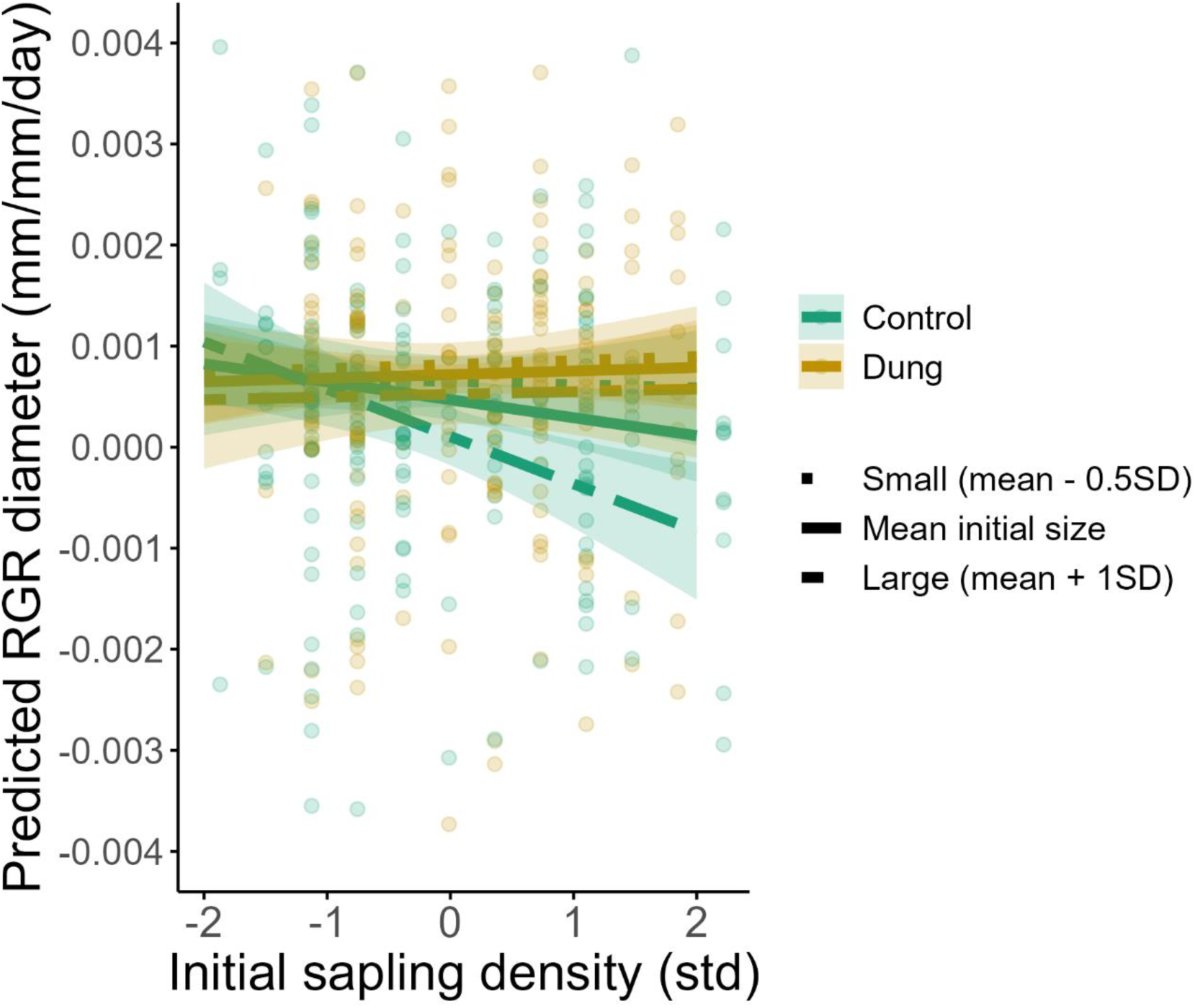
Size-dependent effect of neighborhood competition on sapling growth (RGR_diameter_) under dung treatment and control conditions, with raw data points shown. 17 data points were omitted due to y-axis truncation for better visualization. The buffering provided by dung input against negative density dependence for larger saplings is evident in contrast with control saplings. Estimates derived from the GLMM using *ggpredict*. Small sapling diameter is specified as mean – 0.5 SD initial diameter instead of mean – 1SD which is non-existent in our dataset due to its right skew.

**Figure S3:**
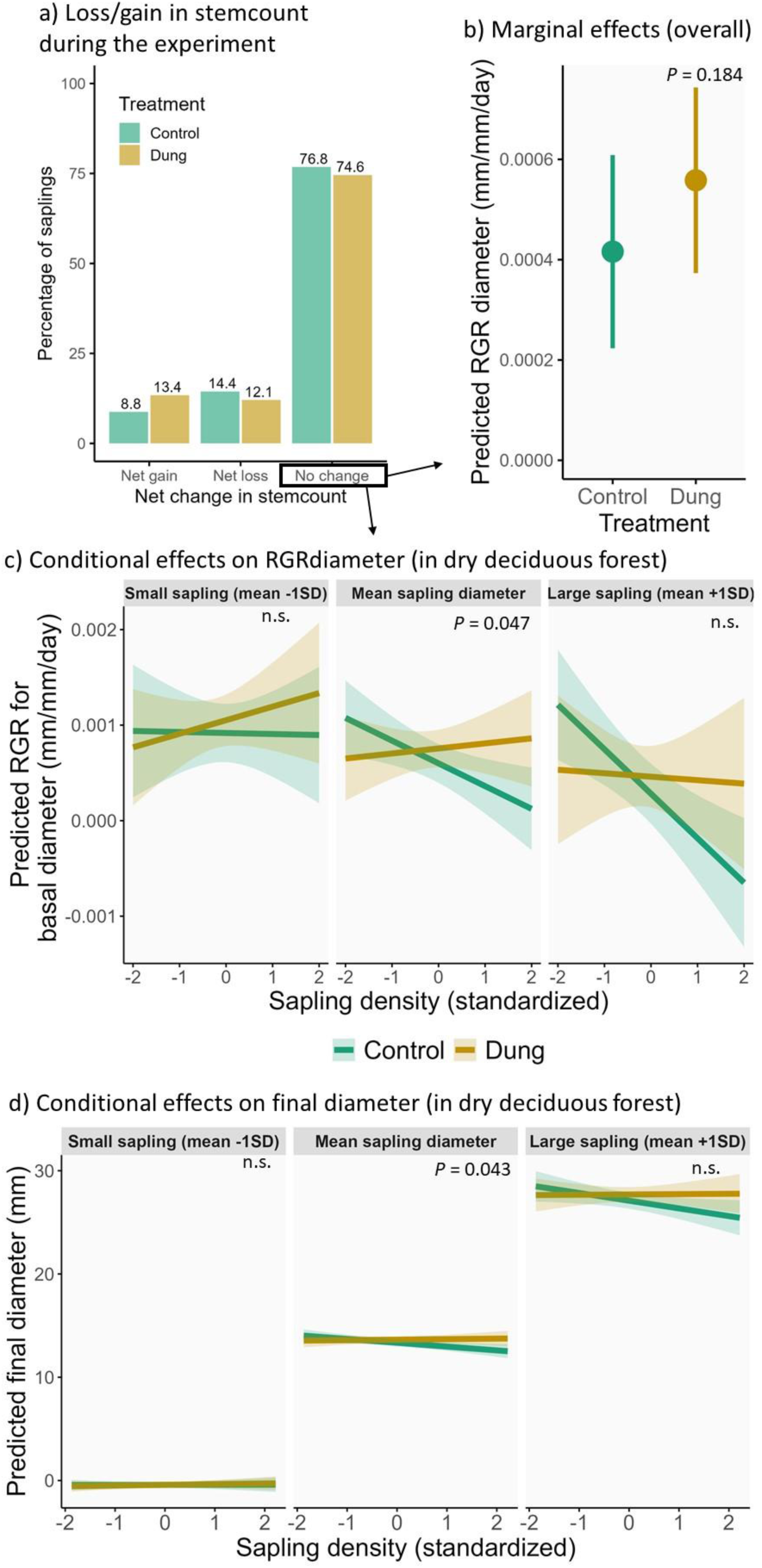
a) Net change in the stem-count of individuals, b) marginal effects and c) conditional effects of dung treatment on RGR_diameter_, for a subset of 316 individuals that did not show any change in stem-count. While the overall growth enhancing effect of dung was statistically not significant (*P*=0.063), the conditional effects (for the deciduous forest which dominated the sample) show that dung treatment significantly buffered the saplings against the negative density-dependent effectsstem-count. This buffering was statistically significant for average-sized saplings (density x treatment interaction) but not larger saplings. d) Similar trends were observed for final basal diameter. See Supplement text A for more details.

**Figure S4:**
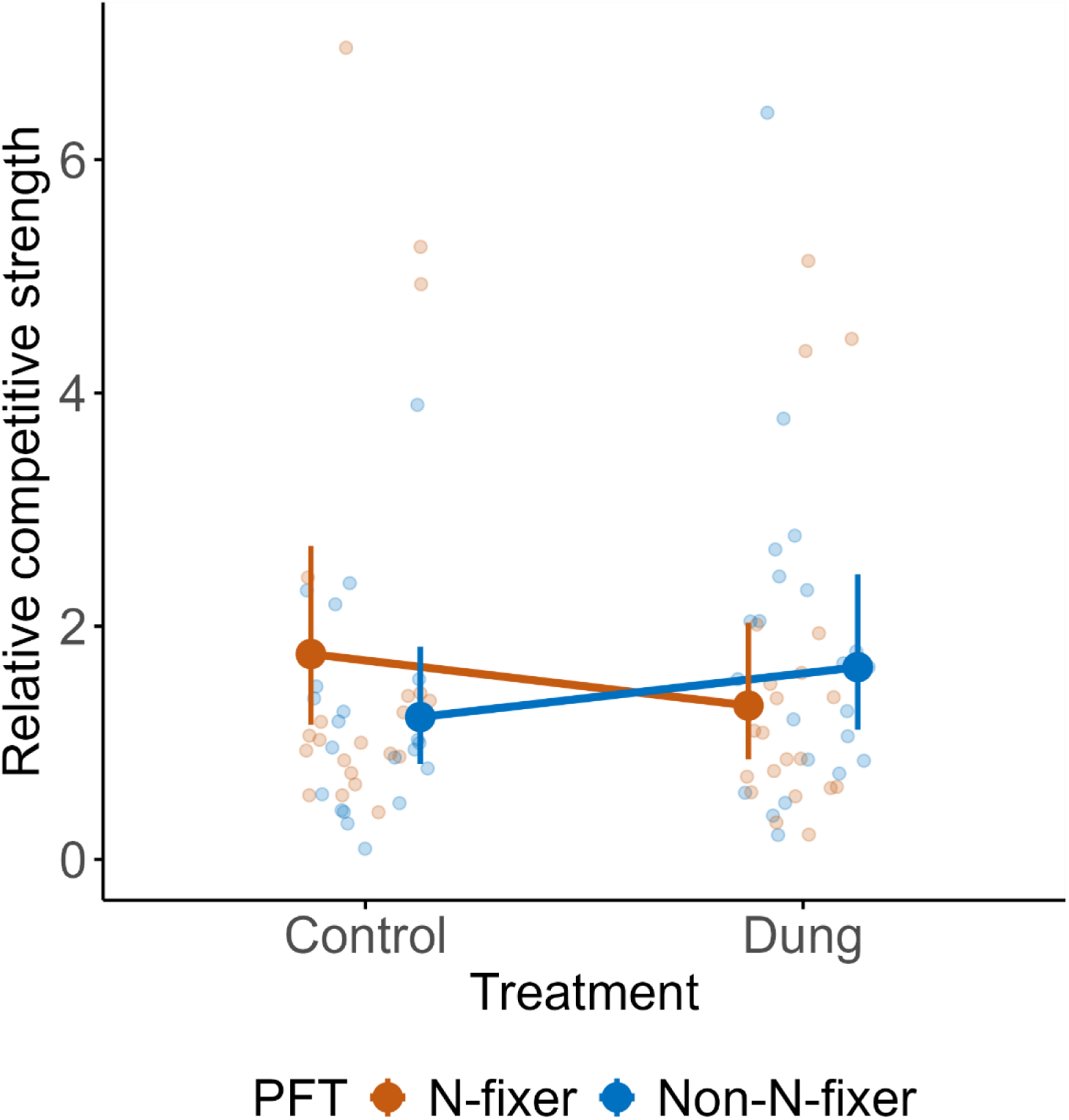
Predicted relative competitive strength (RCS, based on *emmeans*) for N-fixers and non-N-fixers under dung treatment and control in the mesocosm experiment: RCS is significantly >1.0 for N-fixers under control and for non-N-fixers under dung treatment. RCS = biomass under inter-PFT competition / intraspecific competition. Error bars represent 95% confidence intervals around estimated marginal means. This figure is an extension of Figure 4b in the main text, with only raw data points added. One data point for N-fixers has been omitted from control group for better visualization.

**Table S1:** Summary table from a GLMM (with variance structure) of RGR_diameter_ (mm/mm/day) showing conditional effects in dry deciduous forest. Terms of interest are marked blue and significant *P* values are marked bold.

| Conditional effects | Estimate | S.E. | CI lower | CI upper | Pr(>\|z\|) |
| --- | --- | --- | --- | --- | --- |
| **Intercept** (ref: control, DDF, mean sapling size and density) | 0.00047 | 0.00010 | 0.00027 | 0.00067 | **0.000** |
| **Treatment (Dung)** | 0.00025 | 0.00012 | 0.00001 | 0.00049 | **0.040** |
| **Sapling density** | -0.00018 | 0.00010 | -0.00037 | 0.00002 | 0.072 |
| **Initial diameter** | -0.00036 | 0.00012 | -0.00060 | -0.00013 | **0.003** |
| **Forest type – MDF** | -0.00006 | 0.00015 | -0.00035 | 0.00023 | 0.677 |
| **Forest type – Teak** | -0.00031 | 0.00021 | -0.00072 | 0.00010 | 0.143 |
| **Treatment (Dung) × Sapling density** | 0.00021 | 0.00014 | -0.00006 | 0.00048 | 0.120 |
| **Treatment (Dung) × Initial diameter** | 0.00017 | 0.00015 | -0.00013 | 0.00047 | 0.267 |
| **Sapling density × Initial diameter** | -0.00029 | 0.00014 | -0.00056 | -0.00002 | **0.033** |
| **Treatment (Dung) × Sapling density × Initial diameter** | 0.00028 | 0.00018 | -0.00008 | 0.00064 | 0.123 |

**Table S2:**
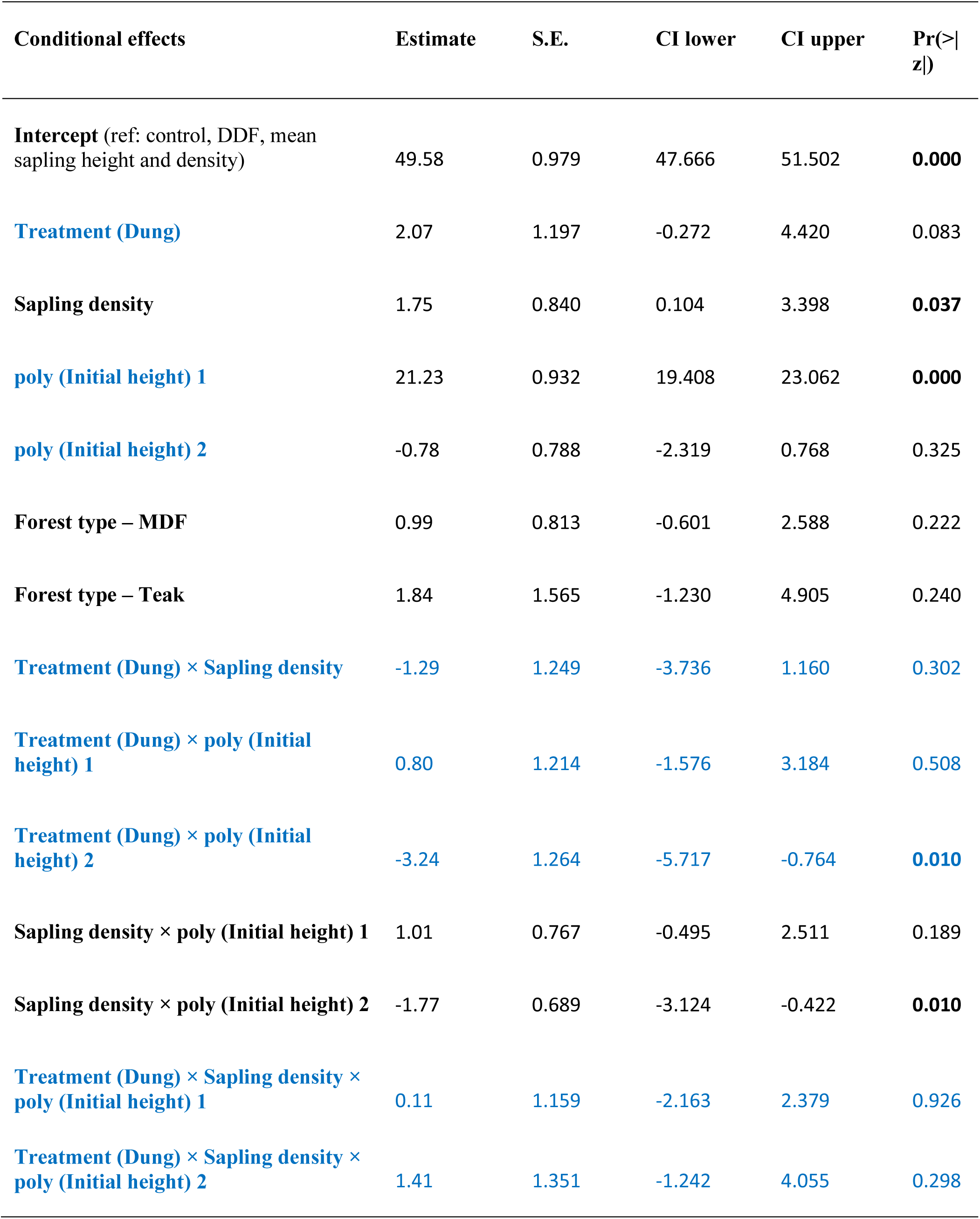
Conditional effects from GLMM showing final height being not significantly increased by dung input. Terms of interest are marked blue and significant effects marked bold (based on 95% CIs).

**Table S3:** Results based on the final sapling basal diameter (mm) of the subset of 316 saplings whose stem-count remained the same. Conditional effects from a GLMM (Student t family of error distribution, with variance structure: dispersion ∼ std. initial diameter) showing dung input buffers saplings of average size against the negative effects of density, for saplings in the dry deciduous forest (which dominated the sample). The marginal positive effect of dung treatment is not significant (*P*=0.157, see Fig S3). Terms of interest are marked blue and significant *P* values are marked bold.

| Conditional effects | Estimate | S.E. | CI lower | CI upper | Pr(>\|z\|) |
| --- | --- | --- | --- | --- | --- |
| **Intercept** (ref: control, DDF, mean sapling size and density) | 13.34 | 0.163 | 13.020 | 13.658 | 0.000 |
| **Treatment (Dung)** | 0.30 | 0.205 | -0.100 | 0.704 | 0.140 |
| **Sapling density** | -0.37 | 0.137 | -0.637 | -0.100 | **0.007** |
| **Initial girth** | 13.76 | 0.248 | 13.272 | 14.246 | **0.000** |
| **Forest type – MDF** | -0.12 | 0.115 | -0.351 | 0.101 | 0.280 |
| **Forest type – Teak** | -0.83 | 0.238 | -1.297 | -0.365 | 0.000 |
| **Treatment (Dung) × Sapling density** | 0.42 | 0.206 | 0.013 | 0.822 | **0.043** |
| **Treatment (Dung) × Initial girth** | 0.30 | 0.332 | -0.349 | 0.955 | 0.362 |
| **Sapling density × Initial girth** | -0.38 | 0.224 | -0.816 | 0.061 | 0.091 |
| **Treatment (Dung) × Sapling density × Initial girth** | 0.36 | 0.334 | -0.295 | 1.015 | 0.282 |

**Table S4:**
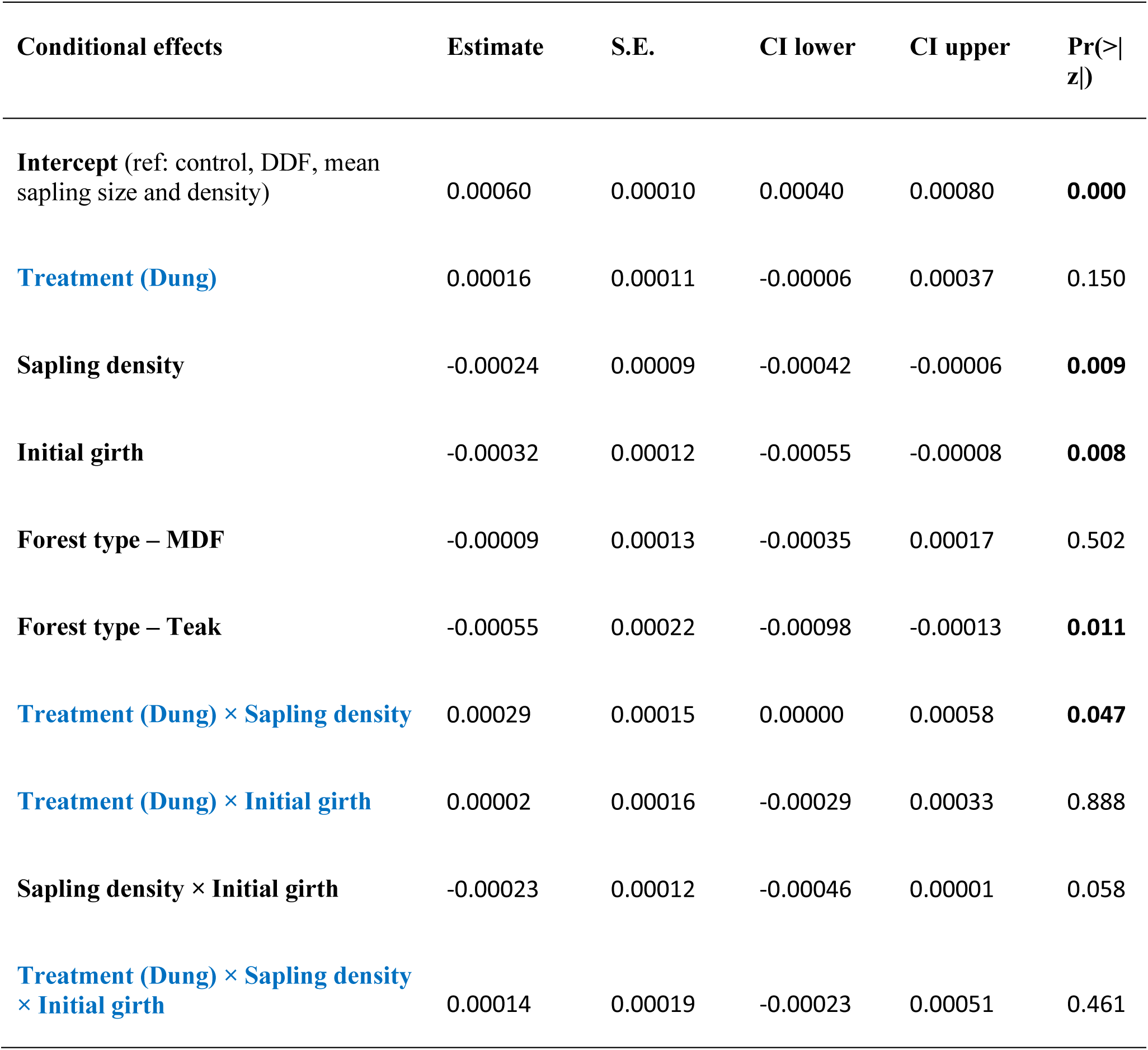
Results based on RGR_diameter_ of the subset of 316 saplings whose stem-count remained the same. Conditional effects from a GLMM (Student t family of error distribution) showing dung input buffers saplings of average size against the negative effects of density, for saplings in the dry deciduous forest (which dominated the sample). The marginal positive effect of dung treatment is not significant (based on *emmeans*, *P*=0.184, see Fig S3). Terms of interest are marked blue and significant *P* values are marked bold.

| Conditional effects | Estimate | S.E. | CI lower | CI upper | Pr(>\|z\|) |
| --- | --- | --- | --- | --- | --- |
| **Intercept** (ref: control, DDF, mean sapling size and density) | 0.00060 | 0.00010 | 0.00040 | 0.00080 | **0.000** |
| **Treatment (Dung)** | 0.00016 | 0.00011 | -0.00006 | 0.00037 | 0.150 |
| **Sapling density** | -0.00024 | 0.00009 | -0.00042 | -0.00006 | **0.009** |
| **Initial girth** | -0.00032 | 0.00012 | -0.00055 | -0.00008 | **0.008** |
| **Forest type – MDF** | -0.00009 | 0.00013 | -0.00035 | 0.00017 | 0.502 |
| **Forest type – Teak** | -0.00055 | 0.00022 | -0.00098 | -0.00013 | **0.011** |
| **Treatment (Dung) × Sapling density** | 0.00029 | 0.00015 | 0.00000 | 0.00058 | **0.047** |
| **Treatment (Dung) × Initial girth** | 0.00002 | 0.00016 | -0.00029 | 0.00033 | 0.888 |
| **Sapling density × Initial girth** | -0.00023 | 0.00012 | -0.00046 | 0.00001 | 0.058 |
| **Treatment (Dung) × Sapling density × Initial girth** | 0.00014 | 0.00019 | -0.00023 | 0.00051 | 0.461 |

**Table S5:** Conditional effects from the GLMM of total sapling biomass under control and dung treatment, and under monoculture and mixture (i.e., inter-PFT) competition in the mesocosm experiment. Terms of *a priori* interest are marked blue and effects different from 0 are marked bold.

|  | Estimate | SE | Lower CI | Upper CI |  |
| --- | --- | --- | --- | --- | --- |
| Intercept (reference: control, N-fixer, monoculture) | 7.614 | 1.604 | 4.470 | 10.758 | **0.000** |
| Treatment (Dung) | -0.325 | 0.513 | -1.331 | 0.680 | 0.526 |
| Competition type (Mix) | 0.434 | 0.751 | -1.038 | 1.906 | 0.564 |
| PFT (Non-N-fixer) | 5.420 | 2.703 | 0.121 | 10.718 | **0.045** |
| Initial size index (standardized) | 2.177 | 0.492 | 1.213 | 3.142 | **0.000** |
| Treatment (Dung) × Competition type (Mix) | -0.075 | 1.164 | -2.358 | 2.207 | 0.948 |
| Treatment (Dung) × PFT (Non-N-fixer) | 0.515 | 2.630 | -4.639 | 5.670 | 0.845 |
| PFT (Non-N-fixer) × Competition type (Mix) | -0.592 | 3.455 | -7.364 | 6.180 | 0.864 |
| Treatment (Dung) × Competition type (Mix) × PFT (Non-N-fixer) | 9.225 | 5.876 | -2.291 | 20.741 | 0.116 |

**Table S6:**
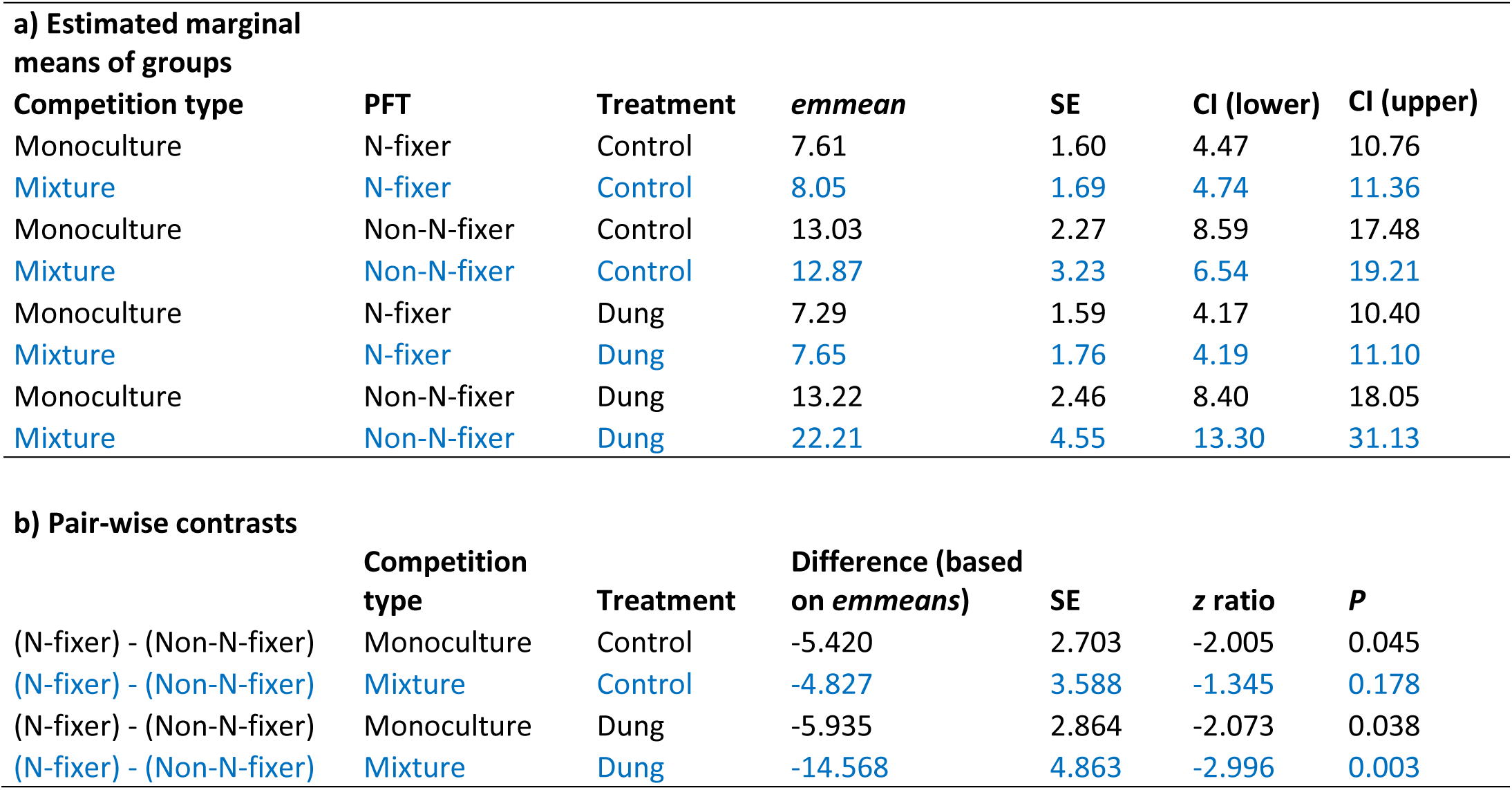
a) Estimated marginal mean sapling biomass (g) of all groups in the mesocosm experiment. b) Pair-wise comparisons for contrasts between plant-functional types within each competition type. Groups/contrasts of interest are marked with blue text.

| **a) Estimated marginal means of groups** |  |  |  |  |  |  |
| --- | --- | --- | --- | --- | --- | --- |
| **Competition type** | **PFT** | **Treatment** | ***emmean*** | **SE** | **CI (lower)** | **CI (upper)** |
| Monoculture | N-fixer | Control | 7.61 | 1.60 | 4.47 | 10.76 |
| Mixture | N-fixer | Control | 8.05 | 1.69 | 4.74 | 11.36 |
| Monoculture | Non-N-fixer | Control | 13.03 | 2.27 | 8.59 | 17.48 |
| Mixture | Non-N-fixer | Control | 12.87 | 3.23 | 6.54 | 19.21 |
| Monoculture | N-fixer | Dung | 7.29 | 1.59 | 4.17 | 10.40 |
| Mixture | N-fixer | Dung | 7.65 | 1.76 | 4.19 | 11.10 |
| Monoculture | Non-N-fixer | Dung | 13.22 | 2.46 | 8.40 | 18.05 |
| Mixture | Non-N-fixer | Dung | 22.21 | 4.55 | 13.30 | 31.13 |

| **b) Pair-wise contrasts** |  |  |  |  |  |  |
| --- | --- | --- | --- | --- | --- | --- |
|  | **Competition type** | **Treatment** | **Difference (based on *emmeans*)** | **SE** | **z ratio** | **P** |
| (N-fixer) - (Non-N-fixer) | Monoculture | Control | -5.420 | 2.703 | -2.005 | 0.045 |
| (N-fixer) - (Non-N-fixer) | Mixture | Control | -4.827 | 3.588 | -1.345 | 0.178 |
| (N-fixer) - (Non-N-fixer) | Monoculture | Dung | -5.935 | 2.864 | -2.073 | 0.038 |
| (N-fixer) - (Non-N-fixer) | Mixture | Dung | -14.568 | 4.863 | -2.996 | 0.003 |

**Table S7:**
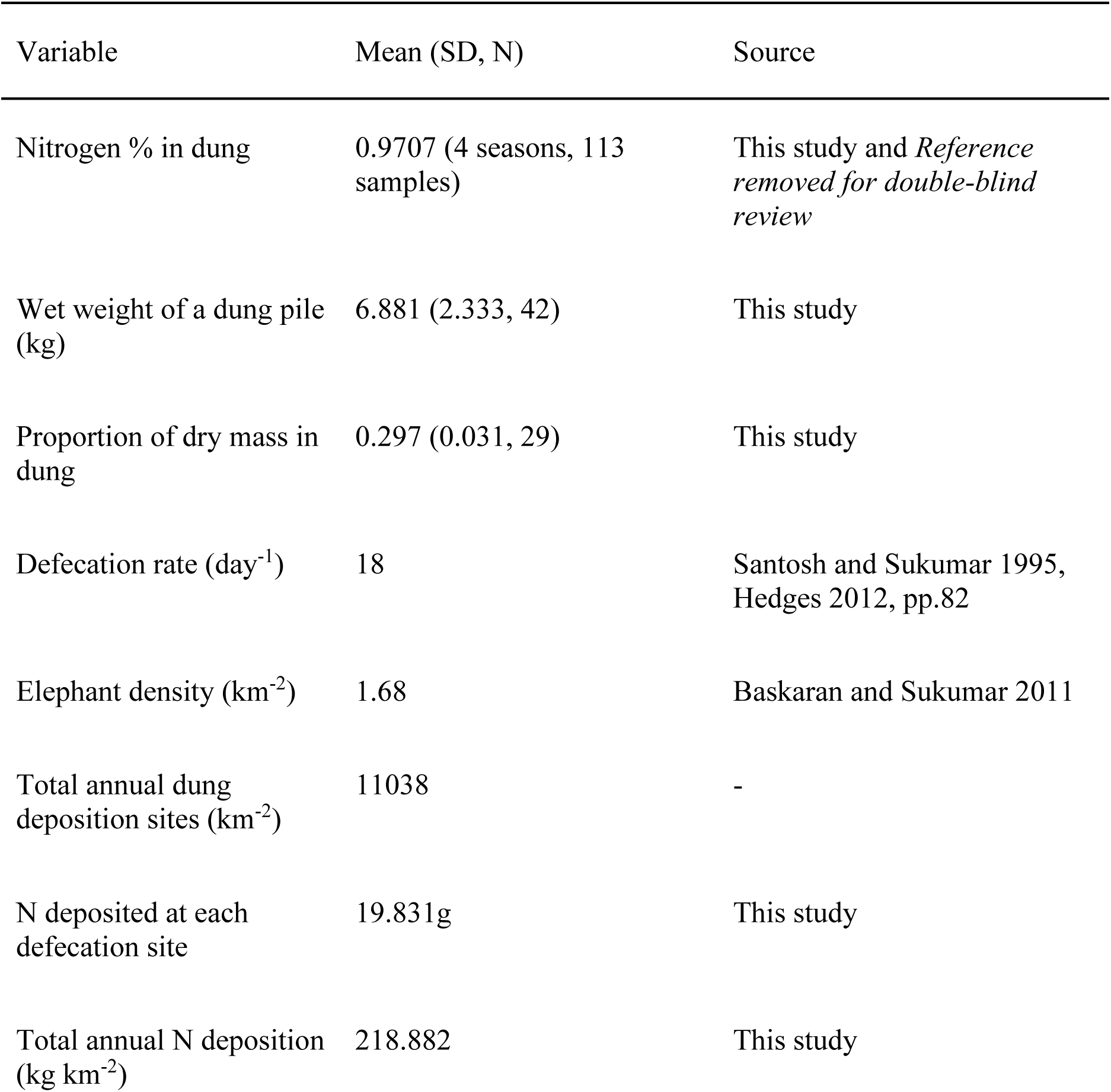
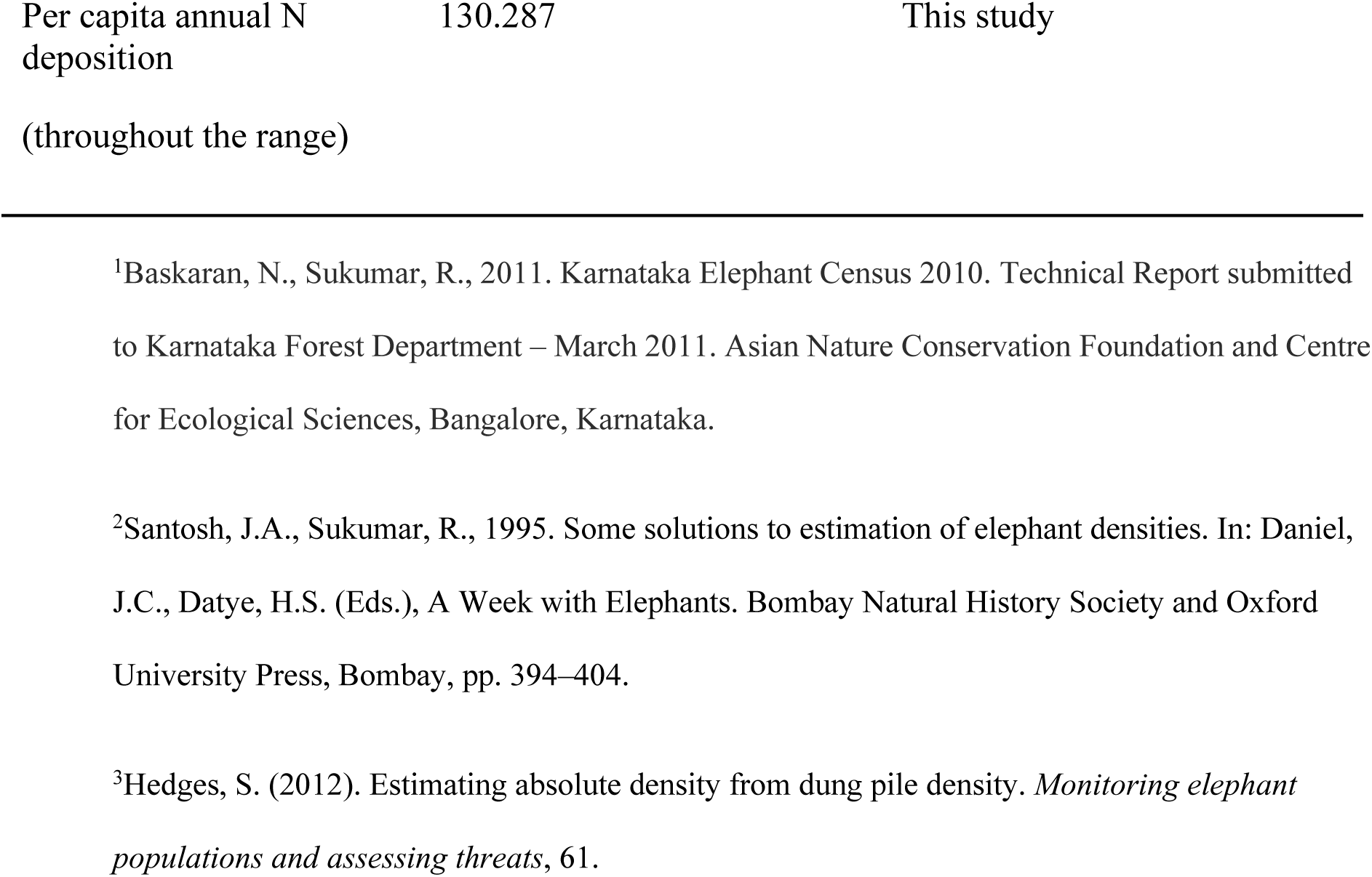
Quantification of annual nitrogen redistribution via dung deposition by elephants in Nagarahole National Park, southern India. N% was averaged from N% quantified at every 3 months: October 2022 (1.151%), January 2023 (1.027%), April 2023 (0.738) based on Gautam et al. 2024, and August 2023 (0.967%) quantified in this study. Estimates of elephant density (∼1.68/km^2^) in Nagarahole were obtained from the Karnataka Elephant Survey 2010 (Baskaran and Sukumar 2011^1^). We used 18/day as the daily defecation rate as reported for Mudumalai in this landscape (Santosh and Sukumar 1995^2^) as well as another study with a large sample size (Hedges 2012 pp. 82^3^).

### Supplement text: A

#### Using predicted RGR_diameter_ to derive percentage and absolute change in diameter

As RGR takes very small values and thus may appear less intuitive, we used RGR_diameter_ to derive two intuitive variables, i.e., percentage change and absolute change in diameter, for different contrasts of treatment, sapling size and density. This conversion is based on following logic.

*a) Deriving absolute change in diameter from predicted RGR_diameter_*

The RGR’s formula, RGR_diameter_ = [ln (final diameter) – ln (initial diameter)]/ *t*

can be rearranged as:

ln (final diameter) = ln (initial diameter) + RGR_diameter_ × *t*,

Both sides can be exponentiated to get:

Final diameter = Initial diameter × e^RGRdiameter^ ^×^ *^t^*

Absolute change in diameter = Final diameter - Initial diameter,

which can be derived from above as

Absolute change in diameter = Initial diameter × (e^RGRdiameter^ ^×^ *^t^*− 1)

We used a constant *t* = 215 days for all size contrasts as it represented the full duration of our field experiment.

*b) Deriving percentage change in diameter from predicted RGR_diameter_*

From the rearrangement,

Final diameter = Initial diameter × e^RGRdiameter^ ^×^ *^t^*,

we can derive relative change as:

(Final diameter - Initial diameter)/ Initial diameter = (e^RGRdiameter^ ^×^ *^t^*− 1),

And thus, percentage change in diameter = 100 × (e^RGRdiameter^ ^×^ *^t^*− 1).

We present in Fig 3 in main text these two derived measures for three discrete densities: mean density, mean – 1SD and mean + 1SD.

#### Effect of dung treatment on net loss and gain in stem-count

We also investigated if the dung-mediated changes in the total diameter and RGR_diameter_, as presented in the main text, came largely from the changes in stem-count of individuals. Such changes in the initial and final stem-count that can arise, for example, from a) natural sprouting/growth trajectory where new stems sprout and grow, and b) partial mortality i.e., mortality of some of the stems. We first quantified the net change in the stem-count of individuals under control and dung treatment for the dataset analyzed in the maintext. Figure S3 presents the % of individuals under each change class. 75% or more individuals experienced no change in stem-count under both control (n=149) and dung treatment (n=167). Compared to control treatment, more individuals experienced net gain in stem-count under dung treatment (30 vs. 17) while fewer individuals showed a net loss in stem-count (27 vs 28). There was no statistical difference in the frequency distributions (Chi squared = 2.499, *P*=0.287, *df*=2).

Since even these minute changes in stem-counts could have contributed to the overall trends presented in the main text, we further used GLMMs (similar to main text) to analyze final size (total diameter) and RGR_diameter_ for only those individuals who showed no change in stem-count (316 individuals). Our findings were largely consistent with results based on the full dataset, as presented in the main text. In the case of final basal diameter, the control individuals were more negatively impacted by sapling density whereas dung treatment provided a buffer against these negative density-dependent effects, althoug this buffering was statistically significant for only average-sized saplings (Figure S3). Similarly, RGR_diameter_ of control individuals was more sensitive to density than the individuals receiving dung input. However, this effect of dung treatment was statistically significant only for average-sized individuals but not for larger individuals, although a trend for buffering was visually evident for large saplings (Figure S3). See supplement Table S3 and Table S4 for more details.

### Supplement text: B

We characterized the climatic space of Nagarahole National Park by extracting temperature data from ERA 5 Land Daily Aggregates (temperature_2m, temperature_2m_min, and temperature_2m_max) and monthly total rainfall data from CHIRPS (total precipitation). These values were calculated for each pixel within Nagarahole National Park and then a mean across all pixels was obtained to capture average conditions. The seasonality of climate is presented in Fig 2 in the main text, whereas annual variation is provided in Table S8 below.

**Table S8:** Inter-annual variation in temperature, rainfall and primary productivity of vegetation measured by normalized difference vegetation index during 2003-2023 in Nagarahole National Park. The years overlapping with this study are marked in grey.

| Year | Mean temp (°C) | Min temp (°C) | Max temp (°C) | Total annual rainfall (mm) |
| --- | --- | --- | --- | --- |
| 2003 | 22.63 | 12.16 | 34.28 | 1621.57 |
| 2004 | 22.35 | 12.63 | 35.47 | 1893.71 |
| 2005 | 22.51 | 12.00 | 34.78 | 2274.65 |
| 2006 | 22.33 | 12.55 | 34.08 | 1909.18 |
| 2007 | 22.35 | 11.77 | 35.67 | 2255.60 |
| 2008 | 22.15 | 12.09 | 34.27 | 1839.26 |
| 2009 | 22.57 | 11.57 | 34.95 | 1912.78 |
| 2010 | 22.66 | 12.31 | 35.72 | 2301.46 |
| 2011 | 22.07 | 11.88 | 33.12 | 1901.65 |
| 2012 | 22.52 | 11.31 | 34.75 | 1748.98 |
| 2013 | 22.41 | 12.53 | 34.96 | 1407.30 |
| 2014 | 22.56 | 13.29 | 35.23 | 1525.38 |
| 2015 | 22.59 | 12.15 | 34.02 | 1454.92 |
| 2016 | 23.19 | 13.55 | 35.80 | 1004.71 |
| 2017 | 22.74 | 13.12 | 35.22 | 1663.78 |
| 2018 | 22.62 | 12.27 | 33.95 | 1776.06 |
| 2019 | 22.85 | 12.20 | 35.34 | 1619.18 |
| 2020 | 22.86 | 13.58 | 34.92 | 1692.06 |
| 2021 | 22.30 | 12.86 | 34.38 | 1857.65 |
| 2022 | 22.41 | 13.08 | 34.50 | 1957.72 |
| 2023 | 22.98 | 11.62 | 35.47 | 1145.62 |

